# A modular Bayesian framework for inferring transmission networks from polyclonal infections, with application to *Plasmodium falciparum*

**DOI:** 10.64898/2026.05.14.725082

**Authors:** Maxwell Murphy, Rasmus Nielsen, T. Alex Perkins, Bryan Greenhouse

## Abstract

**Motivation:** Molecular surveillance and infectious disease transmission network reconstruction can provide compelling evidence for estimating public-health quantities that are difficult to observe directly, including importation, source-sink structure, and differences in onward transmission across locations or intervention strata. These quantities can be expressed as functions of the underlying transmission network, but individual transmission events are rarely observed and many networks may be consistent with the same data. Existing transmission network reconstruction methods leveraging genetic data are often built for settings in which each infection has one dominant source, one representative haplotype, and mutation-driven genetic divergence along transmission chains. These assumptions are poorly matched to polyclonal infections, in which hosts carry multiple genetically distinct clones and recipient infections may reflect contributions from multiple sources. Such infections are common in malaria, tuberculosis, HIV, and many parasitic infections. Methods are needed that can accommodate these data.

**Results:** We present a modular Bayesian framework for estimating directed transmission on sampled cases, where an infection may have no sampled parent, one parent, or several parents, including sources outside the observed panel. Pathogen-specific modules supply likelihoods over candidate parent sets and connect to shared inference that yields marginal directed edge probabilities, posterior mean out-degree, and inclusion probabilities for unobserved parents. We demonstrate our framework with Plasmotrack, a transmission network model for *Plasmodium falciparum* that uses targeted amplicon sequencing data. We implemented these components with a per-locus allele-mixture transmission likelihood, an amplicon genotyping error model, and data augmentation allowing for unobserved parents. Simulations from a biologically informed generative model, under which the inferential per-locus allele-mixture likelihood is misspecified, showed recovery of aggregate network summaries including mean outdegree and mean unobserved-source inclusion, alongside high precision and recall for detecting directed transmission. Other pathogens can reuse the same modular composition after substituting transmission and observation likelihoods.

**Availability:** The Plasmotrack software and documentation are available at https://github.com/eppicenter/plasmotrack. Source code and example datasets are provided under an open-source license.

## 1 Introduction

Transmission networks connect individual infections to population-level questions about infectious disease control. Operational targets often include the proportion of cases attributable to importation, the expected number of downstream infections generated by a case or stratum, directional connectivity between groups or locations, and source-sink structure across surveillance units. In dense outbreak investigations, individual transmission edges may themselves be the target. In many surveillance settings, however, a single reconstructed graph is too specific; transmission events are partly observed, infections are missed, and many graph configurations can imply similar population-level conclusions. The target is therefore often a posterior distribution over network functionals, not a single reconstructed transmission history.

Existing transmission-inference methods have been developed for many pathogens and data settings. Genetic approaches include phylogenetic or tree-based reconstruction, Bayesian who-infected-whom models, and pairwise genetic similarity methods, with applications to SARS [1], foot-and-mouth disease [2, 3, 4], tuberculosis [5], HIV [6], and COVID-19 [7]. These methods are natural when each infection has one dominant source, can be represented by a single haplotype, and accumulates genetic divergence along transmission chains [8]. They are less aligned with pathogens for which hosts routinely carry multiple genetically distinct clones or acquire genetic material from more than one source. In those settings, the relevant compatibility question concerns a candidate set of sources rather than a single parent.

This suggests a separation between the evidence model and network-level inference. Different data sources, including symptom timing, locations, contact histories, genetic data, or combinations of these sources, can all be asked to supply the same object: a nonnegative factor proportional to the conditional probability of each child’s observations given each candidate parent set and the rest of the model. We call each such factor a *parent-set likelihood*. Plasmotrack is a Bayesian framework that uses parent-set likelihoods to infer directed acyclic transmission networks and estimate posterior distributions of network functionals. At the network level, this replaces the requirement that each infection choose one parent with inference over acyclic networks in which an infection may have no sampled parent, one parent, or a set of parents. For complex infections, the parent set is itself an epidemiologically relevant object because it distinguishes single-source infection, multi-source infection, and infection requiring contribution from unsampled or external sources. Posterior marginal probabilities for directed transmission links and for dependence on unsampled or external sources propagate to quantities such as case-level attribution, expected downstream infections, directional connectivity, and the posterior probability that an infection requires an unsampled or external source.

*Plasmodium falciparum* illustrates why this abstraction is useful. Malaria remains a major cause of morbidity and mortality, with an estimated 263 million cases and 597,000 deaths in 2023 [9]. Infections are frequently polyclonal (harboring more than one genetically distinct clone), even in low-transmission settings [10, 11]. Individuals may harbor multiple genetically distinct clones through simultaneous transmission of multiple clones in a single bite (cotransmission), sequential infections over time (superinfection), or both. Unlike mutation-driven phylogenetic signal, malaria genetic variation can be reshuffled during transmission itself, as parasites undergo sexual recombination within the mosquito vector before infecting the next human host. Furthermore, hosts frequently acquire distinct clones from separate vector bites via superinfection. Recipient infections can therefore contain complex genetic mixtures whose compatibility must be assessed jointly across multiple possible sources. These features are incompatible with transmission models that represent each infection as descending from a single recent source. They also make genetic variation epidemiologically informative at the scale of transmission events. Targeted amplicon panels can measure the resulting multi-allelic infections at scale, supporting estimates of importation, source-sink connectivity, and differences in onward transmission across sampled groups.

This paper makes three contributions. First, it defines a parent-set likelihood interface that lets different evidence models score candidate source sets without changing the network-level inference mechanics. Second, it develops Bayesian inference for directed acyclic transmission networks, with posterior summaries for marginal transmission, importation, reproduction, and source-set structure. Third, it instantiates and validates this framework for *P. falciparum*, using an allele-mixture transmission likelihood, an amplicon-sequencing observation model, and simulations that compare Plasmotrack with pairwise genetic approaches across transmission regimes and genetic-diversity settings.

## 2 Methods

We begin with the transmission summaries that motivate the model. We represent the unknown transmission network by an adjacency matrix *A* and define posterior summaries as expectations of functions of *A*, and later extend the representation where needed to include dependence on an unobserved parent. We then introduce the parent-set likelihood model and the topological-ordering parameterization used to estimate these summaries. Finally, we instantiate the framework for *P. falciparum*, summarize the sampler, and describe the simulation study used for validation.

### 2.1 Transmission-Network Summaries

#### 2.1.1 Adjacency-Matrix Representation

Understanding infectious disease transmission dynamics often involves estimating quantities such as the average number of secondary cases per infected individual, the proportion of locally acquired versus imported infections, and differences in onward transmission across sampled groups. These quantities can be expressed as functions of an underlying transmission network. We represent this network by an adjacency matrix *A* ∈ {0, 1}^*n*×*n*^ on *n* infected individuals, where *A*_*ij*_ = 1 if and only if individual *i* transmitted infection to individual *j, A*_*ii*_ = 0, and *A* is constrained to be acyclic. Let A denote the set of all valid acyclic adjacency matrices on these nodes.

Transmission networks are rarely observed directly, so our target estimand is a posterior expectation of a function *f* on adjacency matrices given observed data *X*:

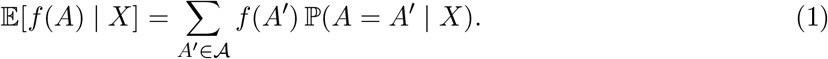

Inference therefore reduces to estimating posterior summaries of *A* that are sufficient for the functionals of interest. Features *f* of practical interest include individual transmission edges, reproduction numbers, importation-related quantities, and source-set summaries. For each *i* ∈ {1, …, *n*}, let *Pa*_*A*_(*i*) = {*j* ∈ {1, …, *n*}: *A*_*ji*_ = 1} denote the parent set of node *i*, i.e., the set of immediate predecessors of *i* in the directed graph defined by *A*.

The following framework treats *X* = (*X*_1_, …, *X*_*n*_) as arbitrary observed data associated with *n* infected individuals. These data may be genetic, epidemiological, spatial, temporal, or any combination of sources, provided the joint model admits a factorization into per-child conditional terms—either directly on the observed data, or at the complete-data level conditional on sampled latent states—that behave as conditional probabilities for each child given its parent set. The *P. falciparum* application developed here uses multi-locus genotypes from polyclonal infections.

#### 2.1.2 Posterior Network Summaries

To accommodate the unobserved-parent construction (supplementary subsection 5.8), we extend *A* to an augmented matrix *Ã* ∈ {0, 1}^(*n*+1)×*n*^ in which index 0 represents the latent unobserved parent: *Ã*_0*j*_ = 1 if and only if the parent set of individual *j* contains an unobserved source. Because transmission-network analyses often ask which entries of *Ã* are likely to be present, the corresponding marginal edge probabilities are also of direct interest. In the model, these probabilities are derived from the primary inferential output: the posterior distribution over augmented parent sets. Let 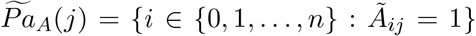 denote the augmented parent set of infection *j*, and let 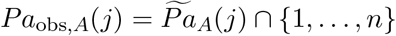 denote its observed parents.

We collect these marginal probabilities in the posterior transmission matrix 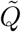:

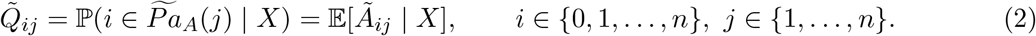

We use 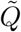 (and its *n × n* submatrix *Q* = 𝔼[*A* | *X*] on observed-infection rows) as a building block for marginal edge and flow summaries. For any linear function *f*, linearity of expectation gives 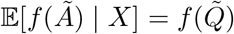, with the analogous identity 𝔼 [*f* (*A*) |*X*] = *f* (*Q*) for summaries involving only observed infections.

Most marginal epidemiological summaries of the transmission network are functionals of 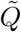. Three are central to the analyses that follow:

- **Posterior edge probability** for the edge *i* → *j*: *Q*_*ij*_, the marginal posterior probability of direct transmission between two observed infections. This is the unit from which network visualizations and edge classifiers are built.
- **Expected out-degree** of node *i*: Σ_*j*≠*i*_ *Q*_*ij*_, the row sum of *Q*, interpretable as an effective reproduction number under appropriate sampling assumptions (see supplementary subsection 5.2).
- **Unobserved-source inclusion probability** for individual 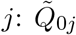, the posterior probability that the parent set of *j* contains a source outside the sampled infections. Averaged across observed infections, 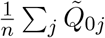 estimates the posterior mean of the realized unobserved-source inclusion proportion.

The augmented unobserved-source indicator should not be interpreted as importation alone. Each true parent contribution is either external to the sampled space, local and observed, or local and missed. The unobserved-source category collapses the external and local-missed contributions, so 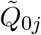 is an upper bound on the posterior probability that infection *j* is imported. More precisely, if *I*_*j*_ is the case-level event that *j* has at least one external parent (i.e., *j* is imported) and *M*_*j*_ is the case-level event that *j* has at least one missed local parent, then 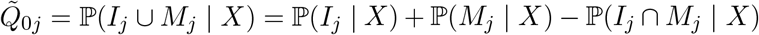. The parent-level states are mutually exclusive, but the case-level events can overlap when an infection has multiple contributing sources. Under a single-source model, the overlap term is zero. Without additional information on local case ascertainment, travel exposure, or external source populations, the model cannot identify the two terms separately. Contrasts in 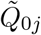 across strata may still be informative when local sampling effort is comparable, because shared missed-local bias partially cancels; supplementary subsection 5.1 gives the corresponding decomposition.

Joint parent-set functionals summarize complementary features of the same posterior distribution that are not determined by 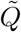 alone. Examples include the probability of multiple contributing sources, such as 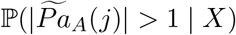 when the unobserved source is counted or ℙ(∣*Pa*_obs,*A*_(*j*) ∣*>* 1 ∣ *X*) when only observed sources are counted, as well as source-set entropy and parent co-inclusion probabilities. The parent-set posterior also decomposes cases into observed-local, mixed observed-and-unobserved, and unobserved-only source categories. These summaries distinguish uncertainty about individual parent identities from uncertainty about the number or type of contributing sources, distinctions that are lost in 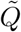 alone.

Each marginal-edge summary is linear in 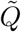, so the MCMC implementation below can estimate uncertainty in the corresponding expected marginal summaries directly from per-step conditional transmission matrices. For example, the MCMC sample 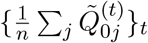 approximates uncertainty in the conditional expectation of the unobserved-source inclusion proportion, and analogous calculations apply to out-degree. Joint source-set summaries are computed from the posterior distribution over complete parent sets, which is available from the same calculation. Supplementary subsection 5.1 gives the mathematical forms used to compute these edge, parent-set, and aggregate summaries from each sampled MCMC state.

#### 2.1.3 Case Sampling and Population-Level Interpretation

The summaries above are defined for the sampled transmission network. Although the augmented unobserved-source category allows parent sets to include sources outside the observed infections, the framework otherwise treats the observed cases as the inference population. In practice, surveillance data are incomplete snapshots of transmission: cases may occur before or after the sampling window, and case collection within the window may be incomplete. These forms of censoring can bias edge-based summaries in different directions depending on the estimand, especially expected out-degree and importation rate. We therefore interpret 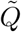-based functionals as summaries of the sampled transmission process unless the sampling assumptions required for population-level interpretation are met. Supplementary subsection 5.2 derives the corresponding censoring consequences for the main summaries.

### 2.2 Parent-Set Likelihood Model

#### 2.2.1 Bayesian Parent-Set Model

Following Huber et al. [12], we apply a Bayesian approach to estimate the posterior distribution of the augmented adjacency matrix *Ã* (subsubsection 2.1.2) and model parameters *θ*:

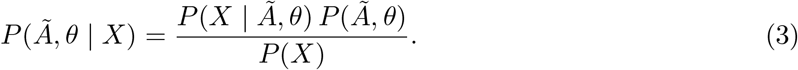

Conditioning on *Ã*, the DAG structure on observed infections factorizes the likelihood into per-child conditional probabilities indexed by the augmented parent set:

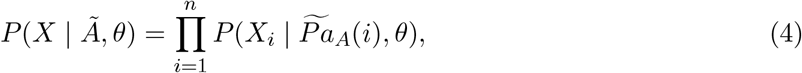

where 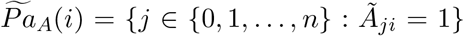 is the augmented parent set of node *i* and index 0 marks inclusion of the latent unobserved source.

For our application to *P. falciparum*, we allow the likelihood to include a latent infection state *Y* to account for imperfect observation of the quantities that inform transmission. The likelihood then becomes:

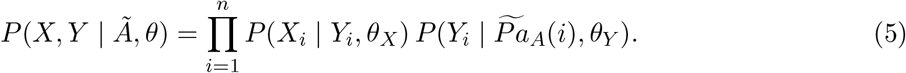

This factorization separates the observation process *P* (*X*_*i*_ ∣*Y*_*i*_, *θ*_*X*_) from the transmission process 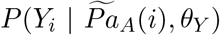, where *Y*_*i*_ is the latent genetic state of infection *i* and *θ*_*Y*_ collects the within-network transmission parameters together with the background-population component that supplies the unobserved-source contribution whenever 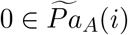 (supplementary subsection 5.8).

The full posterior distribution is:

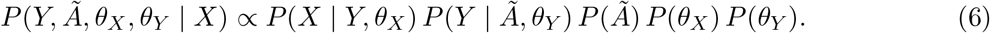

#### 2.2.2 Topological-Ordering Inference

Direct evaluation of the posterior in (6) is computationally intractable because the space of acyclic adjacency matrices grows super-exponentially with the number of nodes. Further, the discrete nature of this space creates highly irregular likelihoods that make standard MCMC sampling inefficient.

Models with latent infection states and unobserved infections as inferred parameters create tight coupling between these parameters and network topology. This coupling makes the likelihood sensitive to parameter changes, requiring simultaneous updates of network topology and other parameters during sampling to maintain reasonable acceptance rates.

We address these challenges using an alternative parameterization based on topological orderings, first applied by Buntine [13] and later popularized by Friedman and Koller [14]. In this framework, we reparameterize our model by the topological ordering ≺, defined as a total ordering over nodes that constrains network topology: if *i < j* in the ordering, then an edge between these nodes, if it exists, must be directed from *i* to *j*.

The ordering ≺ may itself be a model parameter assigned a prior and updated during MCMC, for example a uniform distribution over all possible total orderings of the nodes. Alternatively, ≺ can be induced without a separate ordering prior by any rule that maps auxiliary latent variables to a total order. A natural choice for outbreak data is to rank nodes by inferred infection times: each infection time is obtained from an observation time minus a draw from an infection-to-detection period (IDP) distribution, and the resulting temporal order fixes ≺. We use this induced ordering in the malaria instantiation below; supplementary subsection 5.5 gives the IDP construction and likelihood factorization.

This parameterization enables exact marginalization over the acyclic adjacency matrices compatible with a fixed topological ordering, yielding closed-form expressions for the marginal likelihood *P* (*Y* |≺, *θ*_*Y*_) and for each entry of the conditional edge-probability matrix *Q*^(≺,*Y,θ*)^. It also weakens the coupling between network topology and model parameters, leading to more efficient sampling and improved MCMC mixing. Supplementary subsection 5.3 states the modularity conditions that make this factorization valid and shows how they reduce sums over compatible adjacency matrices to sums over parent sets. Let 𝒰_*j*,≺_ denote the set of candidate parent sets for node *j* compatible ≺ with, with generic element **U**. Under structure modularity, global parameter independence, and parameter modularity,

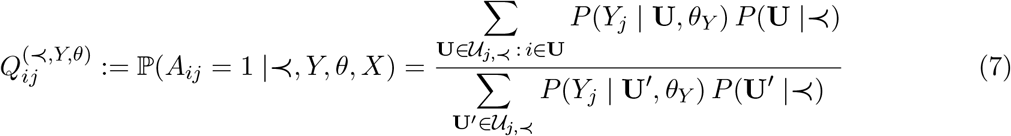

is computable exactly as a sum over parent sets of *j* compatible with ≺ (supplementary Equation 19). The marginal posterior transmission matrix is then

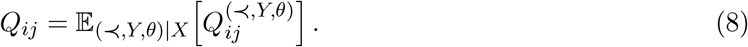

Plasmotrack estimates this expectation by averaging the conditional edge-probability matrices computed at each MCMC state:

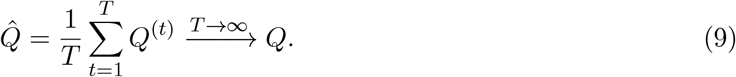

### 2.3 Malaria Instantiation and Validation

#### 2.3.1 Malaria-Specific Likelihoods

The framework developed in subsubsection 2.2.1–subsubsection 2.2.2 is agnostic to the choice of generative model for transmission. In principle, modules can be built from allele-mixture models, spatial or temporal risk models, contact-history models, and application-specific constructions of other forms. For *P. falciparum*, we adopt a per-locus allele-mixture transmission likelihood that captures the salient malaria biology at low computational cost. Supplementary subsection 5.6 gives the full transmission likelihood, supplementary subsection 5.7 gives the observation model, and supplementary subsection 5.8 specifies how unobserved source infections enter the parent-set likelihood.

In this instantiation, *X* is a multi-locus, multi-allelic genotype from each infected individual. This is the natural representation for polyclonal infections, where a single host carries a mixture of genetically distinct clones rather than a single haplotype, and is now routinely produced by targeted amplicon sequencing [15]. Each *X*_*i*_ is a collection of *L* vectors indexed by locus *𝓁*. Each vector *X*_*i𝓁*_ has length *K*_*𝓁*_ and is binary, with *X*_*i𝓁k*_ = 1 indicating the presence of allele *k* and *X*_*i𝓁k*_ = 0 indicating its absence. For example, *X*_*i𝓁*_ may represent a single nucleotide polymorphism (SNP) with length 2, where *X*_*i𝓁*_ = (1, 0) indicates the presence of the first allele and absence of the second. More generally, each *X*_*i𝓁*_ may represent arbitrary genomic data such as microsatellites with *K*_*𝓁*_ possible length polymorphisms or microhaplotypes with *K*_*𝓁*_ possible sequence variants.

The latent state *Y*_*i*_ is the true binary allele-mixture state of infection *i*. The transmission likelihood models bottlenecked sexual recombination in the mosquito vector through a per-locus multinomial in which *m* parasites are sampled from the parent infections and contribute alleles independently across loci; the number *m* is itself a latent variable distributed as a sum of zero-truncated Poissons capturing oocyst-level bottlenecks. To accommodate transmission from sources outside the observed sample, an unobserved parent infection 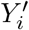 drawn from a background allele-frequency distribution is included alongside any observed parents (supplementary subsection 5.8). The observation likelihood treats per-allele genotyping noise as independent Bernoulli draws around the latent binary allele indicators, with locus-scaled false-positive and false-negative rates appropriate for targeted amplicon sequencing.

These two components plug directly into the framework above, and the inferential procedure in subsubsection 2.2.2 (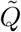 and derived functional summaries) applies unchanged. The components are also interchangeable: a more elaborate transmission likelihood, for example a recombination-graph-based construction that explicitly tracks gametocyte-pairing events at each transmission, or a model that incorporates spatial or temporal covariates of transmission intensity, can be substituted within malaria without modifying any other part of the framework. Substitutions for other infections with mixed within-host genotypes (*Mycobacterium tuberculosis*, HIV, *Trypanosoma, Leishmania*) similarly inherit the rest of the framework.

#### 2.3.2 Sampling and Implementation

The posterior summaries above are estimated from an MCMC trajectory over topological orderings, latent states, and model parameters. We used Metropolis-Hastings updates for continuous parameters with adaptive proposal distributions, specialized samplers for allele frequencies using the self-adjusting logit transform (SALT) [16], and efficient sampling of latent genetic states using both Gibbs and Metropolis-Hastings approaches informed by the transmission network structure. Supplementary subsection 5.10 provides implementation details, parameter initialization, and the sampling algorithms used.

#### 2.3.3 Generative models for simulation

We evaluate the framework using two complementary generative models for the transmission process, chosen to probe behavior under matched and mismatched inferential assumptions. The simplified simulator draws each child latent allele state from a per-locus multinomial over the parent allele mixture, using the same factorization that defines the inferential transmission likelihood (subsubsection 2.3.1); inference is well specified under this simulator. The biologically informed simulator instead walks through the malaria transmission cycle: gametocytes are sampled from each parent, paired gametocytes form oocysts that undergo sexual recombination, and sporozoites are sampled from the resulting oocyst pool and establish in the next host. Each transmitted clone in this simulator therefore descends jointly across loci from a small number of paired-gametocyte oocysts, so child genotypes are coherent whole-parasite mosaics; the inferential per-locus mixture treats loci as exchangeable and cannot impose this joint constraint, so model fits are deliberately misspecified. The biologically informed simulator results are reported as a misspecification stress test, while the simplified-simulator runs reported in the Supplementary Materials serve as the well-specified counterpart. The topology sweep that supplies networks to both simulators is described in subsubsection 2.3.4, and full simulator parameter specifications are in supplementary subsection 5.11.

#### 2.3.4 Simulation Study Design

The simulation study evaluates the malaria-specific likelihood within the general inference framework. We conducted extensive simulations varying network structures and transmission intensities. We explored transmission intensity through network topology parameters including the proportion of founder cases, average number of secondary cases, and proportion of superinfection. We also examined the impact of genetic diversity limitations by varying the complexity of infection (COI) of founder infections and the diversity of genetic loci used for genotyping.

We simulated data using a two-step process: first generating network topologies using a branching process with superinfection merging, then simulating transmission and observation processes. We created 54 distinct network topologies with approximately 200 nodes each by sweeping the parameters used to generate founder status, superinfection, secondary cases, and founder COI. These parameter values were founder probability (0.05, 0.2), superinfection probability (0, 0.05, 0.1), branching-process secondary-case rate (0.5, 1, 1.5), and founder-COI distribution rate (1, 2, 3). For each topology, we generated 10 independent realizations under each of the two generative models described in subsubsection 2.3.3, with three different genotyping panels representing varying levels of genetic diversity.

For a case-censoring stress test, each replicate was additionally analyzed after removing a simple random 50% of infections from the graph (and their incident transmission edges) while retaining genotypes on the remaining nodes and applying the same Dcifer-based parent-set screening as in the fully observed fits. Summaries pair those censored-graph fits to simulation truth computed on the full network before omission (supplementary subsubsection 5.11.6).

Detailed simulation procedures, including parameter specifications, transmission process simulation (a biologically realistic model and a simplified model), observation process simulation, and model fitting settings, are provided in supplementary subsection 5.11. For model fitting, we restricted the parent-set search space using the pre-specified computational constraints described in supplementary subsubsection 5.9.3: each infection could have at most two observed parents plus the augmented unobserved source, and candidate observed parents were filtered using pairwise genetic similarity metrics calculated by Dcifer [17]. In the simulation analyses, candidate observed parents were retained when the lower bound of the 95% confidence interval for pairwise IBD exceeded 0.1.

## 3 Results

We report simulation-based performance for the biologically informed generative model described in subsubsection 2.3.3, under which the inferential per-locus allele-mixture likelihood is misspecified. Unless noted otherwise, fits use the fully observed infection set within each simulation replicate. Figures stratify summaries by founder COI and color points by genotyping panel diversity, because these axes most directly control genetic information available for parent-set discrimination. Additional analyses under a simplified generative model matched to the inferential likelihood and scenario-level diagnostic figures are summarized in subsection 3.5; robustness to random omission of half of infections from the fitted graph is discussed in subsection 3.4.

### 3.1 Edge ranking relative to pairwise IBD

We summarized edge classification performance by integrating precision–recall curves over thresholds applied to posterior marginal edge probabilities and to Dcifer pairwise relatedness scores [17], and by computing the same curve-based summaries for replicate-matched genotype-missing fits that omit the genetic likelihood while retaining the network priors and Dcifer-based parent-set restrictions used in the full model. For each simulation replicate we computed the area under the precision–recall curve (PR-AUC), average precision, and top-*k* precision where *k* was the number of true edges in the simulated network. Posterior scores were evaluated both in the directed sense required for who-infected-whom interpretation and after symmetrizing true edges for an undirected comparator. Dcifer scores are inherently undirected; for those we treated an edge as positive if the true transmission edge existed in either direction, which is optimistic for any use case that requires direction. Genotype-missing baselines were summarized with undirected rankings evaluated against the same undirected notion of truth as symmetrized posterior and Dcifer scores.

Figure 1 aggregates these metrics across replicates, including the genotype-missing undirected series paired to each full-data fit. Posterior directed rankings achieved higher average precision than Dcifer undirected rankings in most founder-COI and genotyping-panel settings, with the largest separation when within-host genetic diversity was high. In the founder COI rate of three and highest diversity panel stratum, replicate medians were 0.54 versus 0.43. For PR-AUC on undirected true edges, symmetrized posterior rankings exceeded Dcifer in the same stratum (0.77 versus 0.69). Pooled across full-network simulation fits, median undirected PR-AUC was 0.75 for Plasmotrack compared with 0.66 for Dcifer, and median undirected average precision was 0.64 compared with 0.43. Genotype-missing undirected rankings were substantially weaker than symmetrized posterior rankings in the same cells, which isolates how much edge ordering is recovered from amplicon genotypes beyond what timing and sparsity priors plus the pairwise similarity screen already imply. At founder COI rate of one, many infections share identical or near-identical genotypes, so all scoring approaches lose resolution, marginal edge probabilities are weak, and PR-type summaries should be interpreted cautiously for ranking individual edges. Posterior undirected summaries track Dcifer more closely in those regimes, as expected when direction is weakly identified. Across panels, Plasmotrack degrades more gently when marker diversity is reduced than Dcifer-based ranking, consistent with borrowing strength across loci and parents under the joint mixture likelihood rather than relying on pairwise IBD point estimates alone.

### 3.2 Recovery of transmission intensity summaries

We next examined scalar summaries derived from the posterior transmission matrix: mean posterior out-degree (average secondary cases among sampled infections when the network is treated as fully observed) and mean posterior probability of inclusion of the augmented unobserved parent, which upper-bounds case-level contribution from outside the sampled set (subsubsection 2.1.2; supplementary subsection 5.1). Figure 2 plots posterior expectations against simulation truth for each replicate.

Posterior mean out-degree tracked the true average number of secondary cases in the sampled network, with modest positive bias that was largest when founder COI was low and panels carried little diversity (for example median bias across replicates of 0.075 at founder COI rate of one with the low-diversity panel versus 0.045 at founder COI three with the high-diversity panel; root mean squared error 0.14 versus 0.089, Table 1), where parent identities were poorly resolved and posterior mass spread across many candidate edges. Mean posterior unobserved-source inclusion slightly exceeded the simulated proportion of infections with an unsampled or external parent in every stratum (mean bias 0.013 to 0.061 on the probability scale, Table 1), with overestimation increasing in founder COI and decreasing with panel diversity (smallest at founder COI rate of one with the high-diversity panel, largest at founder COI three with the low-diversity panel). These patterns illustrate how recovery of the linear 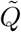 summaries depends on genetic information through parent-set identifiability (subsubsection 2.1.2; supplementary subsection 5.1); distinct biases from incomplete observation of the transmission network itself are summarized in subsection 3.4 alongside the plug-in analysis in supplementary subsection 5.2.

**Supplementary Figure 1:**
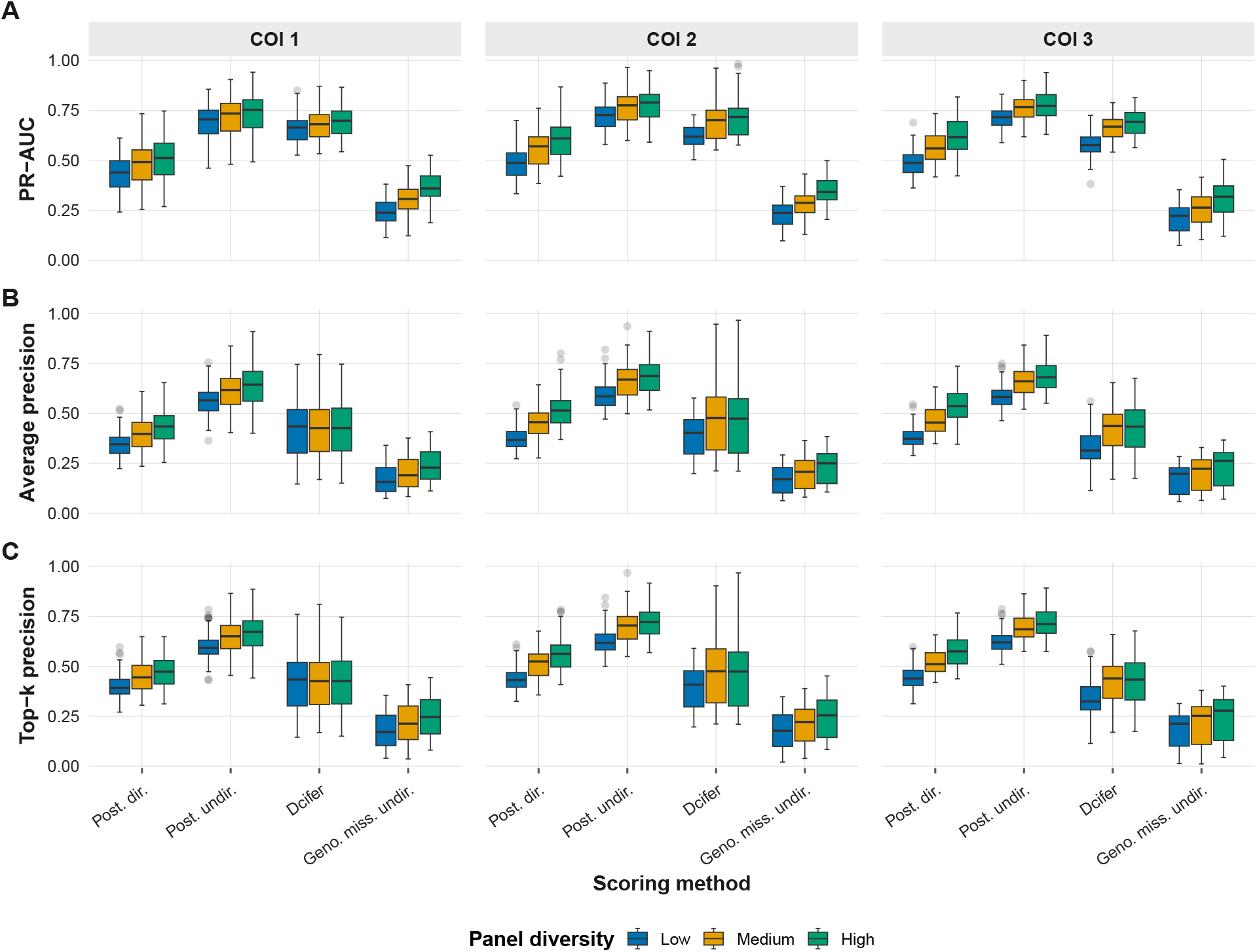
Edge classification metrics under the biologically informed simulator. Each column is a founder COI stratum; colors indicate genotyping panel diversity. Rows show PR-AUC (panel A), average precision (B), and top-*k* precision (C). Within each facet, boxplots contrast four replicate-matched scoring pipelines in fixed horizontal order: posterior directed rankings against directed truth; posterior undirected and Dcifer undirected rankings against symmetrized truth; and genotype-missing undirected rankings from fits that drop the genetic likelihood while retaining the same network priors and Dcifer-based parent-set restrictions.

### 3.3 Parent-set uncertainty with versus without genetic data

Marginal edge metrics alone can understate resolution when posterior mass concentrates on a small set of parent combinations. We therefore compared full posterior fits to fits that retain network priors and Dcifer-based parent-set restrictions but omit the genetic likelihood (genotype missing, or null-genotype runs). For each replicate we report the mean effective parent-set size and the fraction of infections in which the simulated parent token set lies in the nominal 80% marginal credible set over parent sets. Within each founder COI rate by genotyping-panel stratum we summarize replicate-level values by their median across fits.

Figure 3 shows that conditioning on amplicon genotypes sharpens parent-set posteriors relative to the genotype-missing control: lower effective numbers of competing parent sets in every stratum, together with higher 80% credible-set coverage than genotype-missing fits when founder COI and panel diversity were both high. Median effective parent-set size ranged from about 1.5 to 2.1 for the full posterior versus about 10 to 22 for genotype missing. In the founder COI rate of three and highest diversity panel stratum, for example, medians were about 1.6 versus 18 effective sets and 0.79 versus 0.71 for coverage. At founder COI rate of one, median coverages for both fit types clustered near the nominal 0.8 level (about 0.79–0.84 across panels), with only small differences between posterior and genotype-missing fits even though effective parent-set sizes remained much lower for the genotyped posterior. At founder COI rate of three with the low-diversity panel, median posterior coverage was about 0.70, below the nominal 0.8 level. Weak identifiability alone would broaden the posterior over parent sets without displacing mass from the truth, so this shortfall reflects miscalibration in the stratum. The comparison isolates the value of the genetic likelihood on top of the structural and Dcifer-based restrictions already encoded in the model.

**Supplementary Figure 2:**
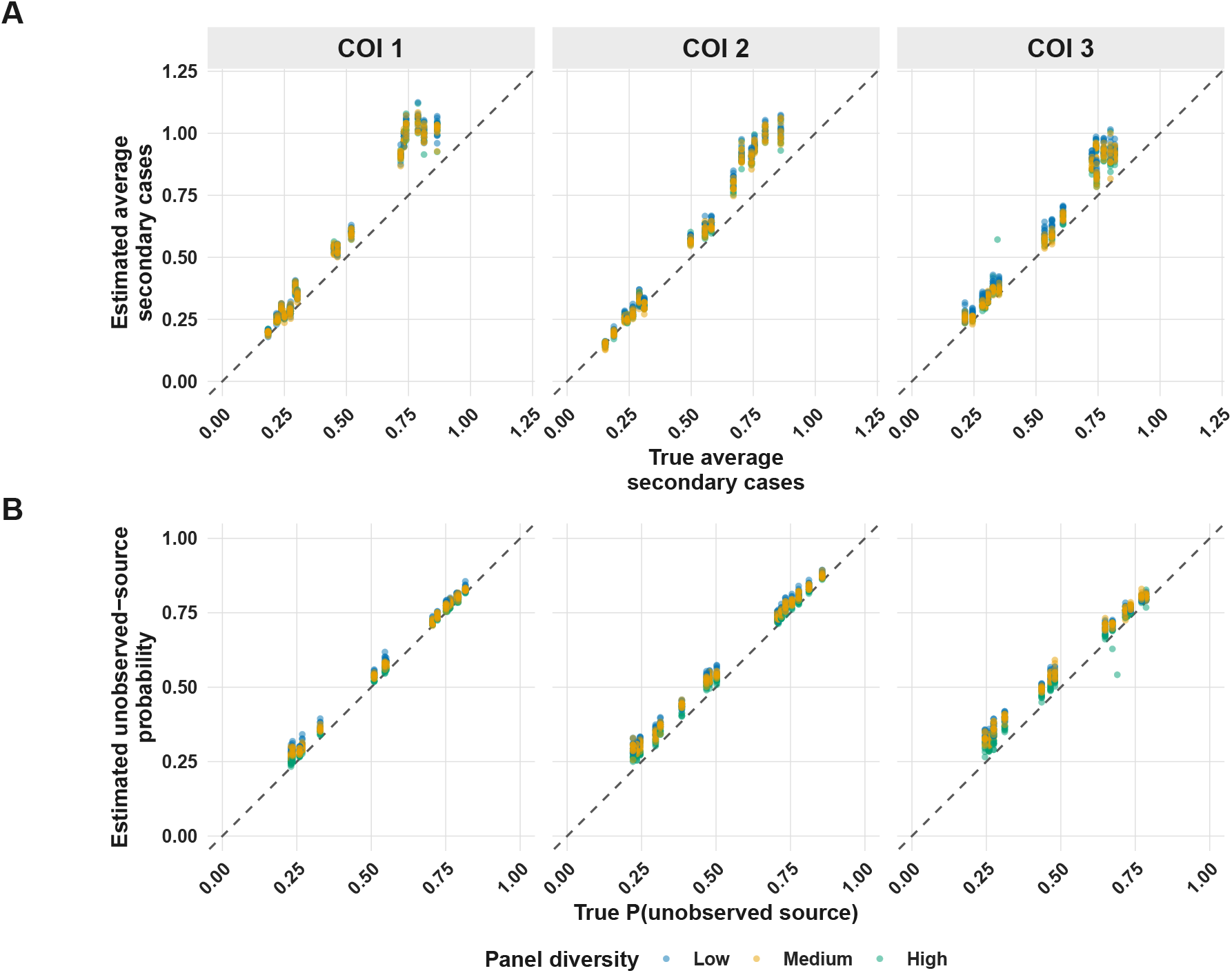
Recovery of average secondary cases and unobserved-source inclusion. Panel A compares true average out-degree in the simulated sampled network to its posterior expectation. Panel B compares the simulated fraction of infections whose parent set includes an unsampled or external source to the mean posterior unobserved-source inclusion probability. Columns stratify by founder COI; colors indicate genotyping panel diversity.

### 3.4 Incomplete observation of cases

We repeated the edge-ranking comparison after fitting the model to the censored 50% infection subset described in subsubsection 2.3.4 (supplementary Figure 7). Replicate medians of undirected PR-AUC for symmetrized posterior scores fell relative to the fully observed fits, with a representative high-information drop from about 0.77 to 0.70 in the founder COI rate of three and highest panel diversity stratum, while directed average precision declined more sharply because many true parents are absent from the fitted graph. Posterior rankings remained comparable to or above Dcifer undirected rankings in most strata, but the margin narrows when censoring removes candidate sources.

**Supplementary Table 1:** Stratum-level error for transmission intensity summaries (biologically informed simulator, fully observed graph). Each row pools posterior fits in one founder COI by genotyping-panel stratum. Out-degree columns summarize recovery of the true average number of secondary cases per analyzed infection: median bias is the median of posterior mean estimate minus truth across fits; root mean squared error (RMSE) is computed on the same replicate-wise differences. Unobserved-source columns summarize recovery of the simulated fraction of infections whose parent set includes an unsampled or external source: mean bias is the mean of posterior mean inclusion probability minus that fraction across fits; RMSE is computed on the same replicate-wise differences. Panel labels low, medium, and high indicate increasing simulated marker diversity.

| Founder COI | Panel | Out-degree |  | Unobserved source |  |
| --- | --- | --- | --- | --- | --- |
|  |  | Med. bias | RMSE | Mean bias | RMSE |
| 1 | Low | 0.075 | 0.142 | 0.032 | 0.036 |
| 1 | Medium | 0.065 | 0.135 | 0.024 | 0.028 |
| 1 | High | 0.069 | 0.137 | 0.013 | 0.016 |
| 2 | Low | 0.059 | 0.116 | 0.051 | 0.055 |
| 2 | Medium | 0.044 | 0.105 | 0.042 | 0.046 |
| 2 | High | 0.046 | 0.104 | 0.026 | 0.029 |
| 3 | Low | 0.075 | 0.111 | 0.061 | 0.067 |
| 3 | Medium | 0.046 | 0.090 | 0.056 | 0.061 |
| 3 | High | 0.045 | 0.089 | 0.033 | 0.040 |

For functional summaries we compared posterior mean out-degree and mean unobserved-source inclusion computed on the censored node set to the same full-network targets used in Figure 2 (supplementary Figure 9). Posterior mean out-degree then systematically underestimates the full-network average number of secondary cases (median replicate bias about −0.08 to −0.10 across founder COI and genotyping-panel strata), whereas mean unobserved-source inclusion is shifted up relative to the full-network fraction of infections with an unsampled or external parent (mean bias about 0.12 to 0.18 on the probability scale across strata). These directions align with using a smaller observed subgraph while retaining truth defined on the complete simulation: local parents are missing from the state space, so marginal transmission intensity on the analyzed nodes falls while probability flows toward the augmented unobserved parent.

Supplementary subsection 5.2 records bias structure for plug-in estimators of network density on induced subgraphs under simple random sampling of nodes and under false positive or false negative edge observation. The posterior summaries here are not plug-in rules and need not coincide with those closed forms, but the same qualitative mechanisms appear when infections are omitted at random: absent parents inflate routes through the unobserved-source category, and edge discriminability weakens when true sources are not in the fitted graph.

The simplified simulator, for which the inferential likelihood is well specified, shows the same qualitative patterns under the same censoring protocol (supplementary Figure 8, supplementary Figure 10), with modestly higher edge-ranking medians and somewhat attenuated functional bias than in the biologically informed runs.

**Supplementary Figure 3:**
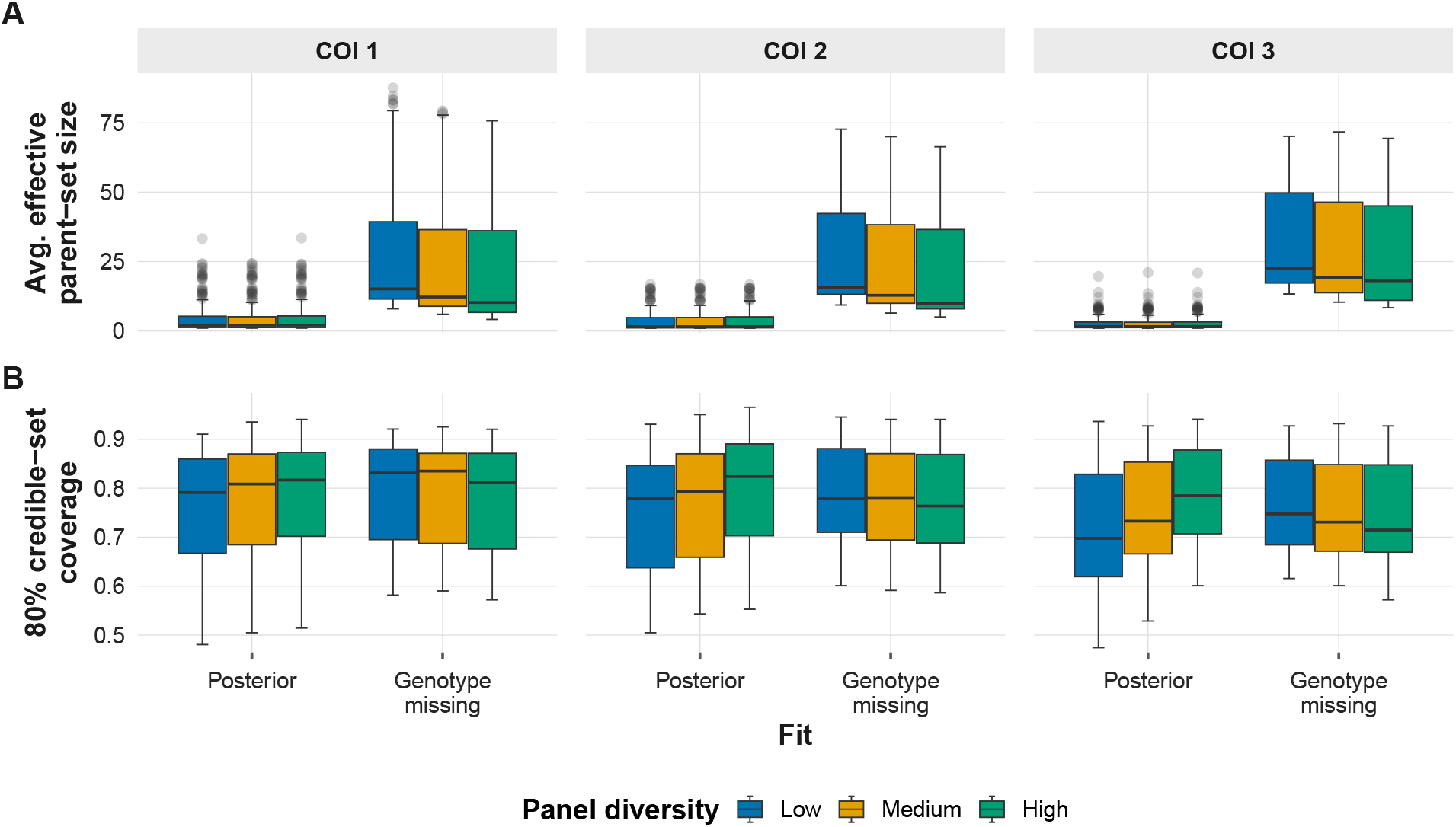
Parent-set sharpness with versus without the genetic likelihood (biologically informed simulator, fully observed graph). Columns are founder COI strata; colors indicate genotyping panel diversity. Rows show mean effective parent-set size (A) and the fraction of infections in which the true parent token set lies in the nominal 80% marginal credible set over parent sets (B). Within each facet, dodged boxplots contrast one full posterior fit against one genotype-missing fit per simulation replicate; genotype-missing fits retain the same network priors and Dcifer-based parent-set restrictions but omit the genetic likelihood.

### 3.5 Supplementary robustness analyses

The Supplementary Materials include multi-panel scenario-group diagnostics (supplementary Figure 5, supplementary Figure 6) that contrast true networks, posterior edge probabilities, an 80% marginal edge credible set, genotype-missing baselines, Dcifer-thresholded graphs, and seed-wise score distributions. Censoring stress tests are interpreted in subsection 3.4. Simplified-simulator tables and figures in the supplement parallel the main-text biologically informed results under full observation, with modestly higher edge-ranking metrics and modestly lower bias and root mean squared error for posterior mean out-degree across COI and panel strata, consistent with the residual biases reported above reflecting biological-model misspecification rather than a fundamental limitation of the framework.

## 4 Discussion

We have developed a general Bayesian framework for inferring transmission networks from modular transmission and observation likelihoods over candidate parent sets, together with a worked genetic instantiation for *P. falciparum*. A single inferential core applies once those likelihoods are specified.

In the simulations above, the *P. falciparum* instantiation recovers transmission structure, direction, and epidemiological functionals across a wide range of intensities and consistently improves on approaches based solely on pairwise genetic similarity.

Malaria combines a large genome, population-level diversity, frequent polyclonal infection, and a mosquito sexual stage that limit plausible likelihood forms while still producing recombination structure that pairwise relatedness underuses. The likelihoods in subsubsection 2.3.1 are tailored to that structure. Within-host genetic diversity is the main control on resolution: at low multiplicity of infection many infections are genetically similar or identical, so directed edge identification is inherently weak.

Genotype-missing fits leave parent sets comparatively diffuse under shared priors and Dcifer-based restrictions (subsection 3.3, Figure 3). Random omission of infections from the fitted graph removes true parents from the state space, which weakens directed rankings more than undirected ones and shifts functional summaries relative to full-network targets (subsection 3.4).

### 4.1 Generality of the framework

Most of what we have presented is independent of the pathogen biology and data source that motivated this work. The Bayesian DAG model (subsubsection 2.2.1), topological-ordering parameterization (subsubsection 2.2.2), Rao-Blackwellized estimator 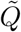 (subsubsection 2.1.2–subsubsection 2.2.2), derived functional summaries (subsubsection 2.1.2), and the censoring properties (supplementary subsection 5.2) hold for any transmission and observation likelihoods that satisfy the structure- and parameter-modularity conditions of the reparameterization.

Within an application, alternative parent-set transmission likelihoods or observation models can be substituted without modifying the shared inferential procedure: the same construction of 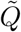 and derived summaries applies once those likelihoods are specified. Richer recombination structure, spatial or temporal transmission covariates, and observation models that encode sampling are natural elaborations of those two components.

Across disease systems, the framework targets the same inferential situation: hosts carry mixtures of distinct clones and clone-level phylogenies are impractical or weak. Examples include *Mycobacterium tuberculosis*, with documented mixed-strain infection and strong bottlenecks but effectively clonal evolution [18]; HIV, with large quasispecies diversity and common superinfection; and kinetoplastids such as *Trypanosoma* and *Leishmania*, for which meiotic recombination and vector transmission resemble *P. falciparum* [19]. Required changes are confined to the parent-set transmission model (for instance, replacing the recombination-and-bottleneck specification of subsubsection 2.3.1 with clonal mixing for TB or HIV) and to the observation model (allele frequencies, microsatellites, or shotgun read data). Uncertainty quantification and behavior under censored observation follow without further structural modification.

### 4.2 Future directions

Several extensions follow naturally from the framework. Transmission modeling could incorporate spatial or temporal covariates of intensity or contact structure, following related person-to-person and reproduction-number formulations for other pathogens [12, 20]. Latent within-host genetics could be extended with explicit recombination and pairing structure along the lines discussed by Camponovo et al. [21], including graph objects related to ancestral recombination graphs. The augmented unobserved parent absorbs unsampled sources, but the observation process does not yet encode which hosts enter the study. Supplementary subsection 5.2 formalizes bias mechanisms for simple subgraph statistics under random node omission and misclassified edges, and subsection 3.4 documents matching qualitative behavior for posterior rankings and functionals when infections are dropped at random. A sampling-aware likelihood that models observation inclusion could in principle correct those biases at fit time rather than only in post hoc comparison to full-network truth.

