## Supplemental methods and figures for "A modular Bayesian framework for inferring transmission networks from polyclonal infections, with application to *Plasmodium falciparum*"

### Supplementary Material for A general Bayesian framework for estimating transmission networks from polyclonal data, with application to *Plasmodium falciparum*

Maxwell Murphy[1], Rasmus Nielsen[2], T. Alex Perkins[3], and Bryan Greenhouse[1]

[1]EPPIcenter program, Division of HIV, ID and Global Medicine, Department of Medicine, University of California, San Francisco, CA, USA
[2]Center for Computational Biology, University of California, Berkeley, Berkeley, CA, USA
[3]Department of Biological Sciences, University of Notre Dame, Notre Dame, IN, USA

2 **Scenario-group network diagnostics (biologically informed simulator).** One representative scenario group (matched simulation factors and genotyping panel tier, ten stochastic realizations; full observation). Top row, left to right: **A**, true directed transmission network among analyzed infections; **B**, posterior marginal directed edge probabilities (edge width and colour encode posterior support); **C**, directed edges in an 80% marginal credible set; **D**, marginal edge probabilities under genotype-missing fits (null genetic likelihood, with network priors and Dcifer-based parent-set restrictions retained); **E**, Dcifer undirected graph retaining pairs whose score exceeds the 0.9 quantile of strictly positive pairwise scores on the representative run. Bottom row: **F**, precision–recall style curves with scoring methods overlaid for the group; **G**, distributions of selected edge or ranking metrics across seeds; **H**, scalar functional summaries aggregated over the same realizations (e.g. average out-degree and unobserved-source summaries). . . . 25
3 **Scenario-group network diagnostics (simplified simulator matched to the inferential likelihood).** Same layout as Figure 2. . . . 26
4 **Edge classification metrics when half of infections are censored from the fitted graph (biologically informed simulator).** Each column is a founder COI stratum; colors indicate genotyping panel diversity. Rows show PR-AUC (panel A), average precision (B), and top-$k$ precision (C). Within each facet, boxplots contrast four replicate-matched scoring pipelines in fixed horizontal order: posterior directed rankings against directed truth; posterior undirected and Dcifer undirected rankings against symmetrized truth; and genotype-missing undirected rankings from fits that drop the genetic likelihood while retaining the same network priors and Dcifer-based parent-set restrictions. Posterior and Dcifer scores are computed on the 50% censored network for each run. . . . 27
5 **Edge classification metrics when half of infections are censored from the fitted graph (simplified simulator).** Same layout as Figure 4. . . . 28
6 **Recovery of average secondary cases and unobserved-source inclusion under 50% case censoring (biologically informed simulator).** Posterior summaries are computed on the 50% censored observed subset for each replicate. Panel A compares true full-network average out-degree to its posterior expectation under censoring. Panel B compares the simulated full-network fraction of infections whose parent set includes an unsampled or external source to the mean posterior unobserved-source inclusion probability under censoring. Columns stratify by founder COI; colors indicate genotyping panel diversity. 29
7 **Recovery of average secondary cases and unobserved-source inclusion under 50% case censoring (simplified simulator).** Same layout as Figure 6. . . . 30

8 **Exact R values for different parent set weighting schemes across a range of network sizes under a fixed constraint on parent sets to include up to 2 observed parents and 1 unobserved parent.** Excluding unobserved infections from possible parent sets results in a higher R value under a fixed topological ordering when weighting all parent sets equally. This situation may arise to some degree when genetic data supports connectivity between infections, but does not differentiate between infections, such as a cluster of genetically identical infections. In contrast, including unobserved infections in possible parent sets results in a lower R value under a fixed topological ordering. This situation would arise when there is insufficient genetic data to differentiate between local transmission and introduction or importation, such as when genetic marker diversity is very low and infections are genetically identical. . . . . . . . . . . . . . . . . . . . . . . . . . 31

### 1 Supplementary Methods

#### 1.1 Derived Quantities: Mathematical Details

Our original problem was to estimate functions or features $f$ of the adjacency matrix $A$ underlying the transmission network (Equation 1). After reparameterizing the model in terms of the topological ordering, we integrate over the acyclic adjacency matrices compatible with each ordering. Let $\mathcal{A}_{\prec}$ denote this set of compatible matrices. While sampling adjacency matrices conditional on the topological ordering is theoretically possible, certain features of interest can be derived directly from the ordering and parent-set weights [1].

We begin by restating the problem of estimating the probability of a feature $f$ under a fixed topological ordering:

$$\mathbb{P}(f \mid \prec, X, Y, \theta_X, \theta_Y) = \frac{P(f, X, Y \mid \prec, \theta_X, \theta_Y)}{P(X, Y \mid \prec, \theta_X, \theta_Y)} \tag{1}$$

where $P(f, X, Y \mid \prec, \theta_X, \theta_Y)$ is the joint probability of feature $f$, the observed data $X$, and the latent genetic state $Y$, obtained by summing over compatible structures containing $f$ weighted by their probability:

$$\begin{aligned} P(f, X, Y \mid \prec, \theta_X, \theta_Y) &= \sum_{A' \in \mathcal{A}_{\prec}} f(A') P(X \mid Y, \theta_X) P(Y \mid A', \theta_Y) P(A' \mid \prec) &(2)\\ &= P(X \mid Y, \theta_X) \sum_{A' \in \mathcal{A}_{\prec}} f(A') P(Y \mid A', \theta_Y) P(A' \mid \prec) &(3) \end{aligned}$$

and $P(X, Y \mid \prec, \theta_X, \theta_Y)$ is the joint probability of the observed data and latent genetic state under a fixed topological ordering:

$$\begin{aligned} P(X, Y \mid \prec, \theta_X, \theta_Y) &= P(X \mid Y, \theta_X) P(Y \mid \prec, \theta_Y) &(4)\\ &= P(X \mid Y, \theta_X) \sum_{A' \in \mathcal{A}_{\prec}} P(Y \mid A', \theta_Y) P(A' \mid \prec) &(5) \end{aligned}$$

The posterior of feature $f$ is thus independent of $X$ and given by:

$$\mathbb{P}(f \mid \prec, Y, \theta_X, \theta_Y) = \frac{\sum_{A' \in \mathcal{A}_{\prec}} f(A') P(Y \mid A', \theta_Y) P(A' \mid \prec)}{\sum_{A' \in \mathcal{A}_{\prec}} P(Y \mid A', \theta_Y) P(A' \mid \prec)} \tag{6}$$

where the numerator's form depends on the specific feature $f$ of interest. The summaries below separate marginal inclusion quantities, which are determined by $\tilde{Q}$, from joint parent-set quantities, which require the full distribution over parent sets.

###### 1.1.1 Conditional Parent-Set Distribution

The basic object is the posterior distribution over parent sets. For a non-augmented parent set $\mathbf{U} \in \mathcal{U}_{i,\prec}$,

$$\mathbb{P}(Pa_A(i) = \mathbf{U} \mid \prec, Y, \theta_X, \theta_Y) = \frac{\sum_{A' \in \mathcal{A}_\prec} \mathbf{1}(Pa_{A'}(i) = \mathbf{U}) P(Y \mid A', \theta_Y) P(A' \mid \prec)}{\sum_{A' \in \mathcal{A}_\prec} P(Y \mid A', \theta_Y) P(A' \mid \prec)} \tag{7}$$

$$= \frac{P(Y_i \mid \mathbf{U}, \theta_Y) P(\mathbf{U} \mid \prec)}{\sum_{\mathbf{U}' \in \mathcal{U}_{i,\prec}} P(Y_i \mid \mathbf{U}', \theta_Y) P(\mathbf{U}' \mid \prec)}. \tag{8}$$

For the augmented model, let $\widetilde{\mathcal{U}}_{i,\prec}$ denote the compatible parent sets after adding the unobserved source indexed by 0. For $\mathbf{U} \in \widetilde{\mathcal{U}}_{i,\prec}$, define

$$w_i(\mathbf{U}) = \frac{P(Y_i \mid \mathbf{U}, \theta_Y) P(\mathbf{U} \mid \prec)}{\sum_{\mathbf{U}' \in \widetilde{\mathcal{U}}_{i,\prec}} P(Y_i \mid \mathbf{U}', \theta_Y) P(\mathbf{U}' \mid \prec)}. \tag{9}$$

This is the conditional posterior probability of parent set $\mathbf{U}$ given the sampled topological ordering, latent genetic states, and model parameters. Let $\mathbf{U}_{\text{obs}} = \mathbf{U} \cap \{1, \ldots, n\}$ denote the observed parents in $\mathbf{U}$.

###### 1.1.2 Marginal Inclusion Summaries

Marginal inclusion summaries depend only on whether each candidate parent appears in the parent set. These are the quantities collected in the posterior transmission matrix $\tilde{Q}$.

For an observed parent $j$ of infection $i$, the conditional edge probability is

$$\mathbb{P}(j \in Pa_A(i) \mid \prec, Y, \theta_X, \theta_Y) = \frac{\sum_{A' \in \mathcal{A}_\prec} \mathbf{1}(j \in Pa_{A'}(i)) P(Y \mid A', \theta_Y) P(A' \mid \prec)}{\sum_{A' \in \mathcal{A}_\prec} P(Y \mid A', \theta_Y) P(A' \mid \prec)} \tag{10}$$

$$= \frac{\sum_{\mathbf{U} \in \mathcal{U}_{i,\prec} : j \in \mathbf{U}} P(Y_i \mid \mathbf{U}, \theta_Y) P(\mathbf{U} \mid \prec)}{\sum_{\mathbf{U}' \in \mathcal{U}_{i,\prec}} P(Y_i \mid \mathbf{U}', \theta_Y) P(\mathbf{U}' \mid \prec)}. \tag{11}$$

The expected out-degree of observed infection $i$ is the sum of its marginal child-inclusion probabilities:

$$\sum_{j=1; j \neq i}^{n} \mathbb{P}(i \in Pa_A(j) \mid \prec, Y, \theta_X, \theta_Y) = \sum_{j=1; j \neq i}^{n} \frac{\sum_{A' \in \mathcal{A}_\prec} \mathbf{1}(i \in Pa_{A'}(j)) P(Y \mid A', \theta_Y) P(A' \mid \prec)}{\sum_{A' \in \mathcal{A}_\prec} P(Y \mid A', \theta_Y) P(A' \mid \prec)} \tag{12}$$

$$= \sum_{j=1; j \neq i}^{n} \frac{\sum_{\mathbf{U} \in \mathcal{U}_{j,\prec} : i \in \mathbf{U}} P(Y_j \mid \mathbf{U}, \theta_Y) P(\mathbf{U} \mid \prec)}{\sum_{\mathbf{U}' \in \mathcal{U}_{j,\prec}} P(Y_j \mid \mathbf{U}', \theta_Y) P(\mathbf{U}' \mid \prec)}. \tag{13}$$

For a child infection $i$, the expected number of observed parents is also a marginal summary:

$$\mathbb{E}[|Pa_{\text{obs},A}(i)| \mid \prec, Y, \theta_Y, X] = \sum_{\mathbf{U} \in \widetilde{\mathcal{U}}_{i,\prec}} |\mathbf{U}_{\text{obs}}| \, w_i(\mathbf{U}). \tag{14}$$

The unobserved-source inclusion probability is

$$\mathbb{P}(0 \in \widetilde{Pa}_A(i) \mid \prec, Y, \theta_Y, X) = \sum_{\mathbf{U} \in \widetilde{\mathcal{U}}_{i,\prec}} \mathbf{1}(0 \in \mathbf{U}) \, w_i(\mathbf{U}). \tag{15}$$

Let $q_i = \mathbb{P}(0 \in \widetilde{Pa}_A(i) \mid \cdot)$ denote this unobserved-source probability. It includes both imported infections, whose parents lie outside the sampled space, and locally transmitted infections whose parents were not sampled. If $I_i$ is the event that infection $i$ is imported and $M_i$ is the event that it has a missed local parent, then $q_i = \mathbb{P}(I_i \cup M_i \mid \cdot)$, so $q_i$ is an upper bound on the importation probability $\mathbb{P}(I_i \mid \cdot)$.

If each infection has a single source and a local source is observed with probability $\rho_i$, then

$$q_i = p_i^{\text{imp}} + (1 - p_i^{\text{imp}})(1 - \rho_i), \tag{16}$$

where $p_i^{\text{imp}} = \mathbb{P}(I_i \mid \cdot)$. For a known, externally estimated, or sensitivity-analysis value of $\rho_i$,

$$p_i^{\text{imp}} = \frac{q_i + \rho_i - 1}{\rho_i}. \tag{17}$$

Even when absolute importation probabilities are not identifiable, contrasts can be useful under comparable sampling. Let $q_s$ be the mean unobserved-source inclusion probability in stratum $s$, and let $p_s^{\text{imp}}$ be the corresponding importation probability. Under the single-source approximation with a common local ascertainment probability $\rho$ across strata,

$$q_a - q_b = \rho \left( p_a^{\text{imp}} - p_b^{\text{imp}} \right). \tag{18}$$

Thus the sign of the unobserved-source contrast matches the sign of the importation contrast, but the magnitude is attenuated by local ascertainment. If ascertainment differs across strata, unobserved-source contrasts remain confounded by missed local transmission and should be interpreted as differences in external-or-unsampled contribution rather than importation alone.

##### 1.1.3 Joint Parent-Set Summaries

Joint parent-set summaries require the full distribution $w_i(\mathbf{U})$ because they depend on combinations of parents within the same source set. The complete parent-set posterior separates three case-level source patterns. These are summaries of the sampled transmission network, not direct population-level importation categories. Under case censoring, missed local infections shift probability toward categories involving the unobserved-source category, so the mixed and unobserved-only categories should be read as evidence for unsampled contribution rather than importation alone:

$$\mathbb{P}(0 \notin \widetilde{Pa}_A(i), |Pa_{\text{obs},A}(i)| > 0 \mid \prec, Y, \theta_Y, X) = \sum_{\mathbf{U} \in \widetilde{\mathcal{U}}_{i,\prec}} \mathbf{1}(0 \notin \mathbf{U}) \mathbf{1}(|\mathbf{U}_{\text{obs}}| > 0)\, w_i(\mathbf{U}), \tag{19}$$

$$\mathbb{P}(0 \in \widetilde{Pa}_A(i), |Pa_{\text{obs},A}(i)| > 0 \mid \prec, Y, \theta_Y, X) = \sum_{\mathbf{U} \in \widetilde{\mathcal{U}}_{i,\prec}} \mathbf{1}(0 \in \mathbf{U}) \mathbf{1}(|\mathbf{U}_{\text{obs}}| > 0)\, w_i(\mathbf{U}), \tag{20}$$

$$\mathbb{P}(0 \in \widetilde{Pa}_A(i), |Pa_{\text{obs},A}(i)| = 0 \mid \prec, Y, \theta_Y, X) = \sum_{\mathbf{U} \in \widetilde{\mathcal{U}}_{i,\prec}} \mathbf{1}(0 \in \mathbf{U}) \mathbf{1}(|\mathbf{U}_{\text{obs}}| = 0)\, w_i(\mathbf{U}). \tag{21}$$

The first term is the probability that infection $i$ is explained by observed local parents only. The second is the probability of mixed observed and unobserved contribution. The third is the probability that no observed parent is included.

Multi-source probabilities measure evidence for more than one contributing source, but their interpretation depends on whether the unobserved-source category is counted. The augmented version counts unobserved-source contribution as a source and is therefore sensitive to censoring:

$$\mathbb{P}(|\widetilde{Pa}_A(i)| > 1 \mid \prec, Y, \theta_Y, X) = \sum_{\mathbf{U} \in \widetilde{\mathcal{U}}_{i,\prec}} \mathbf{1}(|\mathbf{U}| > 1)\, w_i(\mathbf{U}), \tag{22}$$

whereas the observed-only version measures evidence for multiple sampled parents:

$$\mathbb{P}(|Pa_{\text{obs},A}(i)| > 1 \mid \prec, Y, \theta_Y, X) = \sum_{\mathbf{U} \in \widetilde{\mathcal{U}}_{i,\prec}} \mathbf{1}(|\mathbf{U}_{\text{obs}}| > 1)\, w_i(\mathbf{U}). \tag{23}$$

The posterior entropy of the parent-set distribution is a scalar measure of source-set uncertainty, not of biological complexity:

$$H_i = -\sum_{\mathbf{U} \in \widetilde{\mathcal{U}}_{i,\prec}} w_i(\mathbf{U}) \log w_i(\mathbf{U}). \tag{24}$$

High entropy indicates that several parent sets remain plausible after conditioning on the sampled ordering, latent genetic states, and parameters. Co-inclusion probabilities instead measure whether two candidate parents tend to appear together in plausible parent sets:

$$\mathbb{P}(j, k \in \widetilde{Pa}_A(i) \mid \prec, Y, \theta_Y, X) = \sum_{\mathbf{U} \in \widetilde{\mathcal{U}}_{i,\prec}} \mathbf{1}(j \in \mathbf{U})\mathbf{1}(k \in \mathbf{U})\, w_i(\mathbf{U}). \tag{25}$$

##### 1.1.4 Computational Efficiency

These derived quantities can be computed efficiently because they leverage terms already calculated during the MCMC sampling process. Specifically, the probability of the latent genetic state given a parent set, $P(Y_i \mid \mathbf{U}, \theta_Y)$, is computed as part of the likelihood evaluation and can be reused for calculating these derived quantities without additional computational cost.

Interpreting the same marginal summaries under incomplete observation of nodes or edges is less direct if one uses plug-in estimators on an induced subgraph rather than posterior expectations. The next subsection records bias structure for representative plug-in rules.

#### 1.2 Case Censoring

Partially observed networks hinder inference because missing nodes and edges bias plug-in estimators of network statistics, such as the average out-degree [2].

Let $A$ denote the adjacency matrix of the complete directed network without self loops on vertex set $V$, and let $V^* \subseteq V$ be the observed vertices obtained by simple random sampling without replacement. Let $n = |V|$ and $m = |V^*|$. Write $A_{i,j}$ for the directed adjacency indicator from node $i$ to node $j$, and let $I(\cdot)$ denote an indicator.

We define the average out-degree of the complete network as

$$\psi = \frac{1}{n} \sum_{i \in V} \sum_{\substack{j \in V \\ j \neq i}} A_{i,j}. \tag{26}$$

The plug-in estimator based on the sampled vertices is

$$\hat{\psi} = \frac{1}{m} \sum_{i \in V} \sum_{\substack{j \in V \\ j \neq i}} I(i, j \in V^*) A_{i,j} \tag{27}$$

**Proposition 1.1** (Bias of Plug-in Estimator Under Perfect Observation)**.** *Under simple random sampling of $m$ nodes without replacement, the expected value of the plug-in estimator $\hat{\psi}$ is:*

$$\mathbb{E}[\hat{\psi} \mid m] = \frac{m-1}{n-1}\psi \tag{28}$$

*Proof.* Conditional on the number of observed nodes $m$, we have:

$$\begin{aligned}
\mathbb{E}[\hat{\psi} \mid m] &= \mathbb{E}\left[\frac{1}{m}\sum_{i\in V}\sum_{\substack{j\in V\\ j\neq i}} I(i,j\in V^*)A_{i,j}\right] &(29)\\
&= \frac{1}{m}\sum_{i\in V}\sum_{\substack{j\in V\\ j\neq i}} \mathbb{P}(i,j\in V^* \mid m)A_{i,j} &(30)\\
&= \frac{1}{m}\frac{m(m-1)}{n(n-1)}\sum_{i\in V}\sum_{\substack{j\in V\\ j\neq i}} A_{i,j} &(31)\\
&= \frac{m-1}{n-1}\psi &(32)
\end{aligned}$$

where the third equality follows from the fact that under simple random sampling without replacement, $\mathbb{P}(i,j\in V^* \mid m) = \frac{m(m-1)}{n(n-1)}$ for all pairs $(i,j)$ with $i \neq j$. □

The plug-in estimator is attenuated by the conditional inclusion probability of a second node given the first, $\mathbb{P}(j \in V^* \mid i \in V^*, m) = \frac{m-1}{n-1}$: it recovers $\psi$ only up to this factor. This complicates comparisons across settings when observation rates differ.

However, contrasts of average out-degrees between node subgroups remain identified if the sampling mechanism is independent of the subgroup definitions. Let the vertex set $V$ be partitioned into strata $S_1, \ldots, S_K$ based on a node-level attribute, with $n_k = |S_k|$. The true average out-degree for stratum $k$ is defined as

$$\psi_k = \frac{1}{n_k}\sum_{i\in S_k}\sum_{\substack{j\in V\\ j\neq i}} A_{i,j}. \tag{33}$$

The corresponding plug-in estimator is the average out-degree for observed nodes in stratum $k$, calculated within the induced subgraph on $V^*$. Let $m_k = |V^* \cap S_k|$ be the number of observed nodes in stratum $k$. The estimator is

$$\begin{aligned}
\hat{\psi}_k &= \frac{1}{m_k}\sum_{i\in V^*\cap S_k}\sum_{\substack{j\in V^*\\ j\neq i}} A_{i,j} &(34)\\
&= \frac{1}{m_k}\sum_{i\in S_k}\sum_{\substack{j\in V\\ j\neq i}} I(i,j\in V^*)\,A_{i,j}, &(35)
\end{aligned}$$

assuming $m_k > 0$.

We decompose the stratum-$k$ out degree into within and cross-stratum components:

$$\psi_k^{\text{in}} = \frac{1}{n_k} \sum_{i \in S_k} \sum_{\substack{j \in S_k \\ j \neq i}} A_{i,j} \tag{36}$$

$$\psi_k^{\text{cross}} = \frac{1}{n_k} \sum_{i \in S_k} \sum_{\substack{j \in V \\ j \notin S_k \\ j \neq i}} A_{i,j} \tag{37}$$

$$\psi_k = \psi_k^{\text{in}} + \psi_k^{\text{cross}} \tag{38}$$

**Proposition 1.2** (Consistency of Relative Out-degree Estimation)**.** *Under simple random sampling of nodes and the asymptotic regime where $n \to \infty$ with $m/n \to \lambda \in (0,1)$, each stratum proportion $\frac{n_k}{n} \to \pi_k \in (0,1)$, and $n_k \to \infty$, the ratio of stratum-specific out-degree estimators is consistent for the ratio of true stratum-specific out-degrees:*

$$\frac{\hat{\psi}_1}{\hat{\psi}_2} \xrightarrow{p} \frac{\psi_1}{\psi_2} \tag{39}$$

*for any two strata $S_1$ and $S_2$ with $\psi_2 > 0$.*

*Proof.* Under simple random sampling of nodes, the pairwise inclusion probability $\mathbb{P}(i, j \in V^*) = \frac{m(m-1)}{n(n-1)}$ is constant for all pairs $(i, j)$ and thus independent of their stratum membership. For $i \in S_k$:

- if $j \in S_k$ and $j \neq i$, then $\mathbb{P}(i, j \in V^* \mid m, m_k) = \frac{m_k(m_k-1)}{n_k(n_k-1)}$
- if $j \notin S_k$ and $j \neq i$, then $\mathbb{P}(i, j \in V^* \mid m, m_k) = \frac{m_k}{n_k} \cdot \frac{m-m_k}{n-n_k}$

Taking expectations, we have

$$\mathbb{E}[\hat{\psi}_k \mid m, m_k] = \frac{m_k - 1}{n_k - 1} \psi_k^{\text{in}} + \frac{m - m_k}{n - n_k} \psi_k^{\text{cross}} \tag{40}$$

$$\tag{41}$$

In the asymptotic regime where $n \to \infty$ with $m/n \to \lambda \in (0,1)$, each stratum proportion $\frac{n_k}{n} \to \pi_k \in (0,1)$, and $n_k \to \infty$, we have that

$$\frac{m_k - 1}{n_k - 1} \xrightarrow{p} \lambda \tag{42}$$

$$\frac{m - m_k}{n - n_k} \xrightarrow{p} \lambda \tag{43}$$

Assuming bounded average out-degree, we have that

$$\hat{\psi}_k - \mathbb{E}[\hat{\psi}_k \mid m, m_k] \xrightarrow{p} 0 \tag{44}$$

so that

$$\hat{\psi}_k \xrightarrow{p} \lambda \psi_k \tag{45}$$

Therefore, for two strata $S_1$ and $S_2$ with $\psi_2 > 0$ and $m_1, m_2 \to \infty$, we have that

$$\frac{\hat{\psi}_1}{\hat{\psi}_2} \xrightarrow{p} \frac{\lambda\psi_1}{\lambda\psi_2} = \frac{\psi_1}{\psi_2} \tag{46}$$

□

Thus, while the absolute levels of network density are biased by partial observation, the relative densities between subgroups are consistently estimated.

When edges are observed with error, such that the set of observed edges $E^*$ is not a subset of the true edges $E$, inference is further complicated. We consider a common error model where true edges may be missed (false negatives) and non-existent edges may be spuriously observed (false positives). Let $\alpha = \mathbb{P}(A^*_{i,j} = 1 \mid A_{i,j} = 0)$ be the false positive rate, and $\beta = \mathbb{P}(A^*_{i,j} = 0 \mid A_{i,j} = 1)$ be the false negative rate.

The plug-in estimator must now use the potentially erroneous observed edges $A^*_{i,j}$ within the induced subgraph on the sampled nodes $V^*$. Assuming a fixed sample size of $m = |V^*|$, the estimator for the average out-degree is:

$$\hat{\psi}_{\text{est}} = \frac{1}{m} \sum_{i \in V^*} \sum_{\substack{j \in V^* \\ j \neq i}} A^*_{i,j} \tag{47}$$

$$= \frac{1}{m} \sum_{i \in V} \sum_{\substack{j \in V \\ j \neq i}} I(i, j \in V^*) A^*_{i,j} \tag{48}$$

The expected value of this estimator reveals a more complex bias structure than in the case of perfect edge observation.

**Proposition 1.3.** *Under simple random sampling of $m$ nodes without replacement and the specified edge error model, the expected value of the estimator $\hat{\psi}_{est}$ is:*

$$\mathbb{E}[\hat{\psi}_{est}] = \frac{m-1}{n-1} \left[(1-\beta)\psi + \alpha(n-1-\psi)\right] \tag{49}$$

*Proof.* We take the expectation of the estimator, noting that the node sampling and edge error processes are independent:

$$\mathbb{E}[\hat{\psi}_{\text{est}}] = \frac{1}{m} \sum_{i \in V} \sum_{\substack{j \in V \\ j \neq i}} \mathbb{E}[I(i, j \in V^*)] \cdot \mathbb{E}[A^*_{i,j}] \tag{50}$$

The pairwise inclusion probability is $\mathbb{E}[I(i, j \in V^*)] = \frac{m(m-1)}{n(n-1)}$. The expected value of an observed edge is found by the law of total probability:

$$\begin{aligned} \mathbb{E}[A^*_{i,j}] &= \mathbb{P}(A^*_{i,j} = 1 \mid A_{i,j} = 1)\mathbb{P}(A_{i,j} = 1) + \mathbb{P}(A^*_{i,j} = 1 \mid A_{i,j} = 0)\mathbb{P}(A_{i,j} = 0) \\ &= (1-\beta)A_{i,j} + \alpha(1 - A_{i,j}) \end{aligned} \tag{51}$$

Substituting these into the main expression and summing over all $n(n-1)$ possible dyads gives:

$$\mathbb{E}[\hat{\psi}_{\text{est}}] = \frac{m-1}{n(n-1)} \sum_{i \in V} \sum_{\substack{j \in V \\ j \neq i}} [(1-\beta)A_{i,j} + \alpha(1-A_{i,j})] \tag{52}$$

$$= \frac{m-1}{n(n-1)} \left[ (1-\beta) \sum_{i \in V} \sum_{\substack{j \in V \\ j \neq i}} A_{i,j} + \alpha \sum_{i \in V} \sum_{\substack{j \in V \\ j \neq i}} (1-A_{i,j}) \right] \tag{53}$$

$$= \frac{m-1}{n(n-1)} [(1-\beta)|E| + \alpha(n(n-1) - |E|)] \tag{54}$$

$$= (1-\beta)\frac{m-1}{n-1}\frac{|E|}{n} + \alpha\frac{m-1}{n(n-1)}(n(n-1) - |E|) \tag{55}$$

$$= (1-\beta)\frac{m-1}{n-1}\psi + \alpha\frac{m-1}{n-1}\frac{n(n-1) - n\psi}{n} \tag{56}$$

$$= \frac{m-1}{n-1} [(1-\beta)\psi + \alpha(n-1-\psi)] \tag{57}$$

□

The expected value is a sum of the attenuated true signal, $(1-\beta)\psi$, and the additive noise from false positives, $\alpha(n-1-\psi)$, with the entire quantity scaled by the sampling factor $\frac{m-1}{n-1}$. This additive noise term is proportional to the false positive rate and the average number of non-edges per node.

This structure prevents the consistent estimation of relative out-degrees between subgroups. Assume nodes are sampled by simple random sampling without replacement of size $m$, independently of edge misclassification. Let the vertex set $V$ be partitioned into strata $S_1, \ldots, S_K$, for example, based on a node-level attribute, with $n_k = |S_k|$. Let $m_k = |V^* \cap S_k|$ be the number of observed nodes in stratum $k$. Define the stratum-$k$ out-degree parameter $\psi_k$ and its within-stratum and cross-stratum decomposition as $\psi_k^{\text{in}}$ and $\psi_k^{\text{cross}}$ as before. Let the edge misclassification rates be constant across strata.

Define the stratum-$k$ estimator of the average out-degree based on the observed edges $A^*_{i,j}$:

$$\hat{\psi}_k^{\text{est}} = \frac{1}{m_k} \sum_{i \in V^* \cap S_k} \sum_{\substack{j \in V^* \\ j \neq i}} A^*_{i,j} \tag{58}$$

$$\tag{59}$$

**Proposition 1.4.** *Conditional on $m$ and $m_k$, the stratum-$k$ estimator of the average out-degree is:*

$$\mathbb{E}[\hat{\psi}_k^{est} \mid m, m_k] = (1-\beta)\mathbb{E}[\hat{\psi}_k \mid m, m_k] + \alpha((m-1) - \mathbb{E}[\hat{\psi}_k \mid m, m_k]) \tag{60}$$

*Proof.* Given independence of node sampling and edge misclassification, we have that

$$\mathbb{E}[\hat{\psi}_k^{\text{est}} \mid m, m_k] = \frac{1}{m_k} \sum_{i \in S_k} \sum_{\substack{j \in V \\ j \neq i}} \mathbb{P}(i, j \in V^* \mid m, m_k) \mathbb{E}[A^*_{i,j}] \tag{61}$$

$$= \frac{1}{m_k} \sum_{i \in S_k} \sum_{\substack{j \in V \\ j \neq i}} \mathbb{P}(i, j \in V^* \mid m, m_k) \left[(1-\beta) A_{i,j} + \alpha(1 - A_{i,j})\right] \tag{62}$$

$$= \frac{(1-\beta)}{m_k} \sum_{i \in S_k} \sum_{\substack{j \in V \\ j \neq i}} \mathbb{P}(i, j \in V^* \mid m, m_k) A_{i,j} + \frac{\alpha}{m_k} \sum_{i \in S_k} \sum_{\substack{j \in V \\ j \neq i}} \mathbb{P}(i, j \in V^* \mid m, m_k)(1 - A_{i,j}) \tag{63}$$

$$\tag{64}$$

The first term is simply $(1-\beta)$ multiplied by the expectation of the estimator with perfect edge observations, $\mathbb{E}[\hat{\psi}_k \mid m, m_k]$.

The second term is

$$= \frac{\alpha}{m_k} \sum_{i \in S_k} \sum_{\substack{j \in V \\ j \neq i}} \mathbb{P}(i, j \in V^* \mid m, m_k)(1 - A_{i,j}) \tag{65}$$

$$= \alpha \left[ \frac{1}{m_k} \sum_{i \in S_k} \sum_{\substack{j \in V \\ j \neq i}} \mathbb{P}(i, j \in V^* \mid m, m_k) - \frac{1}{m_k} \sum_{i \in S_k} \sum_{\substack{j \in V \\ j \neq i}} \mathbb{P}(i, j \in V^* \mid m, m_k) A_{i,j} \right] \tag{66}$$

$$= \alpha \left[ \frac{1}{m_k} \sum_{i \in S_k} \sum_{\substack{j \in V \\ j \neq i}} \mathbb{P}(i, j \in V^* \mid m, m_k) - \mathbb{E}[\hat{\psi}_k \mid m, m_k] \right] \tag{67}$$

The sum $\sum_{i \in S_k} \sum_{j \in V, j \neq i} \mathbb{P}(i, j \in V^* \mid m, m_k)$ is the expected number of directed pairs $(i, j)$ where $i \in V^* \cap S_k$ and $j \in V^*, j \neq i$, conditional on $m$ and $m_k$. This is simply $m_k(m-1)$.

Therefore:

$$\mathbb{E}[\hat{\psi}_k^{\text{est}} \mid m, m_k] = (1-\beta)\mathbb{E}[\hat{\psi}_k \mid m, m_k] + \alpha \Big( (m-1) - \mathbb{E}[\hat{\psi}_k \mid m, m_k] \Big) \tag{68}$$

□

Rearranging, we have that

$$\mathbb{E}[\hat{\psi}_k^{\text{est}} \mid m, m_k] = (1 - \alpha - \beta)\mathbb{E}[\hat{\psi}_k \mid m, m_k] + \alpha(m-1) \tag{69}$$

demonstrating that the estimator is attenuated by the edge misclassification rates $(1-\alpha-\beta)$ as well as shifted by the false positive rate $(\alpha(m-1))$. Unfortunately, our ratio based identification result no longer holds in general, as the size-dependent bias term introduced by false positives does not cancel. We note, however, that contrasts by differences in average out-degrees may still be detected:

**Proposition 1.5.** *The expected difference between stratum-specific estimators under edge misclassification equals the difference of their perfect-observation expected values, attenuated by the edge misclassification rates:*

$$\mathbb{E}\left[\hat{\psi}_1^{est} - \hat{\psi}_2^{est}\right] = (1-\alpha-\beta)\left(\mathbb{E}[\hat{\psi}_1] - \mathbb{E}[\hat{\psi}_2]\right). \tag{70}$$

*Proof.* By the linearity of expectation, we have:

$$\mathbb{E}[\hat{\psi}_1^{\text{est}} - \hat{\psi}_2^{\text{est}}] = \mathbb{E}[\hat{\psi}_1^{\text{est}}] - \mathbb{E}[\hat{\psi}_2^{\text{est}}] \tag{71}$$

From the previous proposition, we have the expression for the expected value of a single stratum's estimator, which can be written as:

$$\mathbb{E}[\hat{\psi}_k^{\text{est}}] = (1-\alpha-\beta)\mathbb{E}[\hat{\psi}_k] + \alpha(m-1) \tag{72}$$

Substituting this expression for strata $S_1$ and $S_2$ yields:

$$\mathbb{E}[\hat{\psi}_1^{\text{est}} - \hat{\psi}_2^{\text{est}}] = \Big((1-\alpha-\beta)\mathbb{E}[\hat{\psi}_1] + \alpha(m-1)\Big) - \Big((1-\alpha-\beta)\mathbb{E}[\hat{\psi}_2] + \alpha(m-1)\Big) \tag{73}$$

The additive bias term, $\alpha(m-1)$, is common to both expressions. Since we are subtracting, this term cancels out:

$$= (1-\alpha-\beta)\mathbb{E}[\hat{\psi}_1] - (1-\alpha-\beta)\mathbb{E}[\hat{\psi}_2] \tag{74}$$

Factoring out the common term $(1-\alpha-\beta)$ gives the final result:

$$\mathbb{E}[\hat{\psi}_1^{\text{est}} - \hat{\psi}_2^{\text{est}}] = (1-\alpha-\beta)\left(\mathbb{E}[\hat{\psi}_1] - \mathbb{E}[\hat{\psi}_2]\right) \tag{75}$$

$$\tag{76}$$

The additive false-positive shift $\alpha(m-1)$ is common to both strata and cancels, leaving the difference of perfect-observation expectations attenuated by $(1-\alpha-\beta)$. □

This identity is exact in finite samples, but $\mathbb{E}[\hat{\psi}_k \mid m, m_k]$ does not reduce to $\frac{m-1}{n-1}\,\psi_k$ for stratum-specific estimators. From the proof of Theorem 1.2,

$$\mathbb{E}[\hat{\psi}_k \mid m, m_k] = \frac{m_k-1}{n_k-1}\,\psi_k^{\text{in}} + \frac{m-m_k}{n-n_k}\,\psi_k^{\text{cross}}, \tag{77}$$

so the within- and cross-stratum components of $\psi_k$ enter with distinct inclusion weights that depend on the per-stratum sampling fractions. Under the asymptotic regime of Theorem 1.2, both weights converge to $\lambda$ and $\mathbb{E}[\hat{\psi}_k] \to \lambda\psi_k$, so that

$$\mathbb{E}\left[\hat{\psi}_1^{\text{est}} - \hat{\psi}_2^{\text{est}}\right] \longrightarrow (1-\alpha-\beta)\,\lambda\,(\psi_1 - \psi_2)\,. \tag{78}$$

When $\alpha + \beta < 1$, the finite-sample identity already determines the sign of $\mathbb{E}[\hat{\psi}_1^{\text{est}} - \hat{\psi}_2^{\text{est}}]$ from the sign of $\mathbb{E}[\hat{\psi}_1] - \mathbb{E}[\hat{\psi}_2]$. Recovering the sign of $\psi_1 - \psi_2$ requires either this asymptotic regime or additional assumptions on the within- and cross-stratum inclusion weights, for example comparable sampling fractions $m_k/n_k$ across strata together with negligible cross-stratum out-degree.

##### 1.3 Topological ordering parameterization

The topological ordering parameterization addresses the computational challenges of DAG inference by reparameterizing the model in terms of a total ordering over nodes. This approach, first introduced by Buntine [3] and later popularized by Friedman and Koller [1], allows for efficient calculation of marginal probabilities over all acyclic adjacency matrices compatible with a fixed ordering.

Our original model in the main text is given by Equation 6. We can reparameterize this model as:

$$P(Y, \prec, \theta_X, \theta_Y \mid X) \propto P(X \mid Y, \theta_X) P(Y \mid \prec, \theta_Y) P(\prec) P(\theta_X) P(\theta_Y) \tag{79}$$

The observation process $P(X \mid Y, \theta_X)$ remains unchanged. The transmission process $P(Y \mid \prec, \theta_Y)$ is now a function of the topological ordering $\prec$. Under a fixed topological ordering, we can calculate the transmission process by marginalizing over all acyclic adjacency matrices compatible with the ordering:

$$\begin{aligned} P(Y \mid \prec, \theta_Y) &= \sum_{A' \in \mathcal{A}_\prec} P(Y \mid A', \theta_Y) P(A' \mid \prec) && (80) \\ &= \sum_{A' \in \mathcal{A}_\prec} \prod_{i=1}^{n} \left[ P(Y_i \mid Pa_{A'}(i), \theta_Y) \right] P(A' \mid \prec) && (81) \end{aligned}$$

where $\mathcal{A}_\prec$ is the set of all acyclic adjacency matrices compatible with the topological ordering $\prec$.

This formulation alone does not provide computational advantages over the original model, as we still need to calculate the likelihood for each adjacency matrix in $\mathcal{A}_\prec$. However, we can achieve substantial computational gains under three key conditions [1]:

1. *Structure modularity*: The prior $P(A \mid \prec)$ decomposes into a product of probabilities for each parent set:

$$P(A \mid \prec) = \prod_{i=1}^{n} P(Pa_A(i) \mid \prec) \tag{82}$$

2. *Global parameter independence*: The transmission process likelihood decomposes into a product of node-specific probabilities:

$$P(Y \mid A, \theta_Y) = \prod_{i=1}^{n} P(Y_i \mid Pa_A(i), A, \theta_Y) \tag{83}$$

3. *Parameter modularity*: For any two adjacency matrices $A$ and $A'$ with identical parent sets $Pa_A(i) = Pa_{A'}(i) = \mathbf{U}$ for node $i$:

$$P(Y_i \mid \mathbf{U}, A, \theta_Y) = P(Y_i \mid \mathbf{U}, A', \theta_Y) \tag{84}$$

Under these conditions, we can rewrite the transmission process likelihood as:

$$P(Y \mid \prec, \theta_Y) = \prod_{i=1}^{n} \sum_{\mathbf{U} \in \mathcal{U}_{i,\prec}} P(Y_i \mid \mathbf{U}, \theta_Y) P(\mathbf{U} \mid \prec) \tag{85}$$

where $\mathcal{U}_{i,\prec}$ is the set of all possible parent sets for node $i$ compatible with the topological ordering $\prec$. This formulation provides substantial computational speedup compared to directly marginalizing over adjacency matrices in $\mathcal{A}_\prec$.

Our model satisfies the global parameter independence and parameter modularity conditions. However, we must carefully address the structure modularity condition through our choice of prior.

#### 1.4 Prior specification and computational implications

Adjacency-matrix priors are often chosen for computational convenience rather than strong theoretical justification. When conditioning on topological ordering, this choice has important implications for the induced prior on network structures.

Consider a uniform prior $P(A \mid \prec)$ over all acyclic adjacency matrices compatible with ordering $\prec$. This prior induces a non-uniform prior $P(A)$ on the overall matrix space. To understand why, consider two extreme cases:

A fully linear network, where each node connects to exactly one parent and one child (except root and leaf nodes), is compatible with only one ordering: the linear ordering of the nodes. In contrast, a fully disconnected network, where every node is a root, is compatible with all possible orderings. Thus, the uniform prior $P(A \mid \prec)$ actually favors sparser network structures [1].

This bias toward sparsity is likely appropriate for transmission networks, which are typically sparsely connected in practice. For our model, we adopt a prior with structure modularity that favors smaller parent sets for each node.

#### 1.5 An informed topological ordering

Our model depends on the topological ordering $\prec$, which we now model using temporal information from real-world outbreak data. Without additional information, we could treat $\prec$ as uniformly distributed over all possible orderings. However, outbreak data typically include observation times that provide valuable constraints on the likely ordering of infections.

Let $T$ denote the collection of observation times, where $T_i$ is the observation time for infection $i$. The time between infection and observation, called the infection-to-detection period (IDP), varies by individual. In malaria, this period frequently depends on symptomatic status: symptomatic individuals typically have shorter IDPs because they seek healthcare earlier, while asymptomatic individuals may remain undetected for extended periods.

Following Huber et al. [4], we model the IDP probability for individual $i$ with symptomatic status $S_i$ as:

$$\mathbb{P}(\tau_i = t) \sim \begin{cases} IDP_S & \text{if } S_i = 1 \\ IDP_A & \text{if } S_i = 0 \end{cases} \tag{86}$$

where $IDP_S$ and $IDP_A$ are the IDP distributions for symptomatic and asymptomatic individuals, respectively. We treat these distributions as known and use the parameterizations provided by Huber et al. [4].

The infection time for individual $i$ is $T_i - \tau_i$. The temporal ordering of infections induces the topological ordering: if $T_i - \tau_i < T_j - \tau_j$, then $i < j$ in $\prec$. This approach constrains the topological ordering without making strong assumptions about transmission timing. We impose no probabilistic constraints on the timing between parent and child infections, only on their relative ordering.

Our complete likelihood becomes:

$$P(Y, \tau, \theta_X, \theta_Y \mid X) \propto P(X \mid Y, \theta_X) P(Y \mid \prec(\tau), \theta_Y) P(\tau) P(\theta_X) P(\theta_Y) \tag{87}$$

where $\prec(\tau)$ is the topological ordering induced by the infection times derived from the IDP collection $\tau$.

#### 1.6 Modeling the transmission process

The malaria parasite transmission process begins when a mosquito feeds on an infected human host, ingesting blood containing potentially multiple genetically distinct gametocytes. Within the mosquito midgut, these gametocytes differentiate into gametes and undergo sexual recombination, producing genetically diverse haploid sporozoites that mature in the mosquito salivary glands. Upon subsequent blood feeding, the infected mosquito may transmit these genetically recombined parasites to a new human host, completing the transmission cycle.

To model this complex biological process mathematically, we develop a simplified framework that captures the key genetic dynamics while remaining computationally tractable. We assume mutation is negligible, so the genetic state of an infection at a given time is a composition of the genetic states of its parent infections. Alleles may be lost due to bottleneck effects during transmission, where only a subset of all genetically recombined parasites pass from mosquito to human.

For each locus, the genetic state of the parent infections defines a probability distribution over the possible sampled alleles in the child infection. The probability of drawing a parasite of genotype $Y_i$ from a collection of parent infections $Pa_A(i)$ is given by

$$P(Y_i \mid Pa_A(i), \theta_Y) = \sum_{m=|Pa_A(i)|}^{S} \prod_{\ell=1}^{L} P(Y_{i\ell} \mid Pa_A(i), m) P(m \mid Pa_A(i), \theta_Y) \tag{88}$$

where $|Pa_A(i)|$ is the number of parent infections, $m$ is the total number of parasites sampled from the parent infections, and $S$ is a predefined maximum number of parasites sampled for numerical purposes.

We model the number of parasites sampled from the parent infections as a latent variable, specifically as a sum of $|Pa_A(i)|$ zero-truncated Poisson random variables with maximum sum limited to $S$ [5]. Conditional on the number of parasites sampled and the genetic state of the parent infections, we model the genetic state of the child infection as independent across genetic loci. At each locus, the genetic state follows a multinomial distribution with allele probabilities parameterized by the parent infections' genetic state.

The probability of allele $k$ at locus $\ell$ given the genetic state of the parent infections $Pa_A(i)$ is the sum of the relative frequency of the allele in the parent infections, normalized by the total number of parent infections:

$$f(k, \ell, Pa_A(i)) = \frac{1}{|Pa_A(i)|} \sum_{Y_{i^*\ell} \in Pa_A(i)} \frac{\mathbb{I}(Y_{i^*\ell k} = 1)}{\sum_{k'=1}^{K_\ell} \mathbb{I}(Y_{i^*\ell k'} = 1)} \tag{89}$$

where $K_\ell$ is the number of alleles at locus $\ell$.

For example, consider a locus with 5 possible alleles and 3 parent infections. If the genetic state of parent infections $Pa_A(i)$ at this locus is represented by the matrix

$$Y_{\cdot\ell} \in Pa_A(i) = \begin{bmatrix} 1 & 0 & 0 & 0 & 0 \\ 0 & 1 & 0 & 1 & 0 \\ 0 & 0 & 0 & 1 & 1 \end{bmatrix} \tag{90}$$

where each row represents the genetic state of a parent infection and each column represents a possible allele, then the derived probability vector is $\left[\frac{1}{3}, \frac{1}{6}, 0, \frac{1}{3}, \frac{1}{6}\right]$.

The probability of the latent genetic state $Y_i$ given the parent set $Pa_A(i)$ and number of strains transmitted $m$ is

$$P(Y_i \mid Pa_A(i), m) = \prod_{\ell=1}^{L} \sum_{y^* \in \mathcal{Y}_{Y_{i\ell}}} \frac{m!}{y_1^*! \cdots y_{K_\ell}^*!} \prod_{k=1}^{K_\ell} f(k, \ell, Pa_A(i))^{y_k^*} \tag{91}$$

where $\mathcal{Y}_{Y_{i\ell}}$ is the set of nonnegative integer count vectors $y^* \in \mathbb{Z}_{\geq 0}^{K_\ell}$ with $\sum_{k=1}^{K_\ell} y_k^* = m$ whose positive-count support exactly matches the binary vector $Y_{i\ell}$, that is, $y_k^* \geq 1$ whenever $Y_{i\ell k} = 1$ and $y_k^* = 0$ whenever $Y_{i\ell k} = 0$.

#### 1.7 Modeling the observation process

We modeled observed genetic data $X_i$ as a function of the latent true genetic state $Y_i$ and error rates $\epsilon_i^+$ and $\epsilon_i^-$, where $\epsilon_i^+$ is the probability of a false positive and $\epsilon_i^-$ is the probability of a false negative for individual $i$. The probability of observing genetic data $X_i$ for individual $i$ conditional on the latent true genetic state $Y_i$ and error rates $\epsilon_i^+$ and $\epsilon_i^-$ is given by:

$$P(X_i \mid Y_i, \epsilon_i^+, \epsilon_i^-) = \prod_{\ell=1}^{L} \prod_{k=1}^{K_\ell} \begin{cases} \frac{\epsilon_i^+}{K_\ell} & \text{if } X_{i\ell k} = 1 \text{ and } Y_{i\ell k} = 0 \\ 1 - \frac{\epsilon_i^-}{K_\ell} & \text{if } X_{i\ell k} = 1 \text{ and } Y_{i\ell k} = 1 \\ \frac{\epsilon_i^-}{K_\ell} & \text{if } X_{i\ell k} = 0 \text{ and } Y_{i\ell k} = 1 \\ 1 - \frac{\epsilon_i^+}{K_\ell} & \text{if } X_{i\ell k} = 0 \text{ and } Y_{i\ell k} = 0 \end{cases} \tag{92}$$

where $L$ is the number of genetic loci and $K_\ell$ is the number of alleles at locus $\ell$.

#### 1.8 Sources of transmission

Every transmission network contains at least one root node representing an infection that cannot be explained by other observed infections. These root infections arise from two possible sources: locally acquired cases whose parent infections were missed during surveillance, or imported cases from outside the study population.

To account for these unobserved parent infections, we augment our model with a latent unobserved parent infection $Y_i'$ for each observed infection $i$. This latent parent can serve as the source of infection $i$ when no other parent infections are observed, or when the observed parent set is incomplete.

The unobserved-parent indicator therefore identifies contribution from outside the sampled infections, not the mechanism by which that source was absent from the sample. It also does not by itself imply that an infection has no sampled local parent, because the unobserved source can appear in the same parent set as observed infections. Without additional information, the model does not distinguish external importation from missed local transmission.

The unobserved parent infection $Y_i'$ is drawn from a background population characterized by parameters $\theta_{Y'} = (\pi_{Y'}, \lambda_{Y'})$. Here, $\pi_{Y'}$ represents allele frequencies across genetic loci in the background population, and $\lambda_{Y'}$ is the rate parameter of a zero-truncated Poisson distribution governing the multiplicity of infection for unobserved cases. During inference, $Y_i'$ is explicitly instantiated and updated as part of the MCMC state, so the transmission likelihood conditions directly on this sampled binary genotype rather than analytically marginalizing over it.

The transmission process can incorporate both observed and unobserved parents. Let $\widetilde{Pa}_A(i)$ denote the augmented parent set for infection $i$, including the unobserved source when present:

$$P(Y_i \mid \widetilde{Pa}_A(i), \theta_Y, \theta_{Y'}) = \sum_{m=|\widetilde{Pa}_A(i)|}^{S} \prod_{\ell=1}^{L} P(Y_{i\ell} \mid \widetilde{Pa}_A(i), m) P(m \mid \widetilde{Pa}_A(i), \theta_Y) \times \tag{93}$$

$$(\mathbf{1}(Y_i' \notin \widetilde{Pa}_A(i)) + \mathbf{1}(Y_i' \in \widetilde{Pa}_A(i)) P(Y_i' \mid \theta_{Y'})) \tag{94}$$

where $\mathbf{1}$ is the indicator function that is 1 if the condition is true and 0 otherwise.

The likelihood of the unobserved parent infection $Y_i'$ is:

$$P(Y_i' \mid \theta_{Y'}) = \sum_{s=1}^{\infty} P(s \mid \lambda_{Y'}) \prod_{\ell=1}^{L} \sum_{y^* \in \mathcal{Y}_{Y'_{i\ell}}} \frac{s!}{y_1^*! \cdots y_{K_\ell}^*!} \prod_{k=1}^{K_\ell} \pi_{\ell k}^{y_k^*} \tag{95}$$

#### 1.9 Priors and prespecified model constraints

To complete our Bayesian model specification, we specified prior distributions for all unknown parameters and incorporated external epidemiological knowledge through model constraints. We adopted standard conjugate priors where possible to facilitate efficient sampling, and organized our specification into three components: model parameter priors, epidemiological constraints, and computational constraints.

##### 1.9.1 Model parameter priors

For the parameters that governed our model's behavior, we specified the following prior distributions (Supplementary Table 1):

| Parameter | Description | Distribution | Hyperparameters |
|---|---|---|---|
| $\epsilon^+$ | False positive rate | Beta | $(\alpha_{\epsilon^+}, \beta_{\epsilon^+})$ |
| $\epsilon^-$ | False negative rate | Beta | $(\alpha_{\epsilon^-}, \beta_{\epsilon^-})$ |
| $\pi_Y$ | Allele frequencies | Dirichlet | $\alpha = 1$ |
| $\lambda_Y$ | COI rate for source infections | Gamma | $(\alpha_{\lambda_Y}, \beta_{\lambda_Y})$ |
| $\lambda_M$ | Transmission rate from parent set | Gamma | $(\alpha_{\lambda_M}, \beta_{\lambda_M})$ |
| $\beta_U$ | Parent set size parameter | Beta | $(\alpha_{\beta_U}, \beta_{\beta_U})$ |

**Supplementary Table 1:** Prior distributions and hyperparameters for model parameters. All hyperparameters were user-specified based on domain knowledge and preliminary analyses.

##### 1.9.2 Epidemiological constraints

We incorporated external epidemiological knowledge through infection-to-detection period (IDP) distributions, which characterized the time between infection and detection for symptomatic and asymptomatic individuals. These distributions were based on empirical estimates from Huber et al. [4] and were not estimated from our data. The IDP distributions informed the temporal ordering of infections in our transmission network and were shown in Figure 1.

##### 1.9.3 Computational constraints

Although the model reparameterized in terms of topological ordering is computationally more efficient than directly marginalizing over DAGs, evaluating all possible parent sets for a given node remains challenging.

The number of possible parent sets for a node is $\sum_{k=1}^{n} \binom{n}{k} = 2^n - 1$, where $n$ is the total number of possible parent nodes and $k$ is the number of parents in the parent set. This grows exponentially and becomes computationally prohibitive for large collections of nodes.

To address this computational challenge, we applied two prespecified constraints that were biologically reasonable for malaria transmission networks:

1. **Maximum parent constraint**: We limited the number of possible parents to a maximum value deemed reasonable for the given dataset. In malaria transmission settings, an upper bound of $n = 2$ observed parent infections plus an unobserved parent was reasonable and explained the vast majority of observed transmission while allowing for superinfection.

2. **Genetic similarity constraint**: We constrained which nodes could serve as parents for each node based on genetic similarity. Infections directly connected in transmission networks typically had substantial genetic similarity, as low similarity would have resulted in low transmission likelihood. We applied an a priori constraint for each node on which other nodes could be considered as possible parents using metrics such as genetic distance or identity by descent measures. Transmission networks were typically sparsely connected, so the number of possible parents for a given infection was generally small compared to the complete set of observed infections.

These constraints provided substantial computational speedups in model fitting without significantly impacting results, as they reflected biologically plausible transmission patterns in malaria populations.

#### 1.10 Sampling and Inference

##### 1.10.1 Implementation

Our MCMC algorithm employs efficient computational strategies to handle the complex dependency structure of the model. The implementation uses modular design principles with transparent caching of calculations, ensuring computational feasibility while maintaining flexibility for future model extensions. Our implementation is available at https://github.com/eppicenter/plasmotrack.

##### 1.10.2 Sampling continuous parameters

We update each continuous parameter componentwise with a random-walk Metropolis-Hastings step that conditions on the current latent genetic state $Y$ and topological ordering $\prec$. The relevant target is the conditional posterior

$$P(\theta_X, \theta_Y \mid X, Y, \prec) \propto P(X \mid Y, \theta_X)\, P(Y \mid \prec, \theta_Y)\, P(\theta_X)\, P(\theta_Y), \tag{96}$$

which follows from Equation 79 by holding $Y$ and $\prec$ fixed. The transmission factor $P(Y \mid \prec, \theta_Y)$ is the parent-set sum in Equation 85, so it is computed without any further marginalization over $Y$. Continuous parameters are initialized from their prior distributions, and proposals are generated by a Gaussian random walk centered at the current value with adaptive variance $\sigma_\theta$ tuned during burn-in to a target acceptance rate of 0.234:

$$\epsilon \sim \mathcal{N}(0, \sigma_\theta) \tag{97}$$
$$\theta^{\text{prop}} = \theta + \epsilon. \tag{98}$$

Let $\theta \in \theta_X \cup \theta_Y$ denote the scalar component being updated, and write $(\theta_X^{\text{prop}}, \theta_Y^{\text{prop}})$ for the parameter vector obtained by replacing that component with $\theta^{\text{prop}}$ and leaving every other component at its current value. The proposal is accepted with probability

$$A(\theta, \theta^{\text{prop}}) = \min\left(1,\ \frac{P(X \mid Y, \theta_X^{\text{prop}})\, P(Y \mid \prec, \theta_Y^{\text{prop}})\, P(\theta_X^{\text{prop}})\, P(\theta_Y^{\text{prop}})}{P(X \mid Y, \theta_X)\, P(Y \mid \prec, \theta_Y)\, P(\theta_X)\, P(\theta_Y)} \cdot \frac{g(\theta \mid \theta^{\text{prop}})}{g(\theta^{\text{prop}} \mid \theta)}\right). \tag{99}$$

The Gaussian proposal is symmetric, so the proposal-density ratio cancels. Componentwise updates further simplify the likelihood ratio: when $\theta \in \theta_X$ only the genotyping factor $P(X \mid Y, \theta_X)$ depends on $\theta$ and the transmission factor cancels, and when $\theta \in \theta_Y$ only the transmission factor $P(Y \mid \prec, \theta_Y)$ depends on $\theta$ and the genotyping factor cancels. Writing $L_X(\theta_X) = P(X \mid Y, \theta_X)$ and $L_Y(\theta_Y) = P(Y \mid \prec, \theta_Y)$, the acceptance probability reduces to

$$A(\theta, \theta^{\text{prop}}) = \min\left(1,\ \frac{P(\theta^{\text{prop}})}{P(\theta)} \cdot \begin{cases} L_X(\theta_X^{\text{prop}})/L_X(\theta_X) & \theta \in \theta_X, \\ L_Y(\theta_Y^{\text{prop}})/L_Y(\theta_Y) & \theta \in \theta_Y. \end{cases}\right) \tag{100}$$

Allele frequencies $\pi_Y$ and $\pi_{Y'}$ live on the simplex and are updated with the SALT sampler described below; the acceptance ratio there has the same conditional structure but uses the SALT proposal density in place of the symmetric Gaussian.

##### 1.10.3 Sampling allele frequencies

Allele frequencies are initialized randomly on the unit simplex of dimension $K_\ell$, where $K_\ell$ is the number of alleles at locus $\ell$. We sample allele frequencies using the self-adjusting logit transform (SALT) sampler [6] with Metropolis-Hastings acceptance.

##### 1.10.4 Sampling latent genetic states

Latent genetic states are initialized to the observed genetic states. Unlike most model parameters, the latent genetic state $Y$ is a discrete random variable composed of collections of binary vectors. For a parent set $\mathbf{U}$, write $Y^{\mathbf{U}}$ for the elementwise indicator of allele presence across $\mathbf{U}$, so that $Y^{\mathbf{U}}_{\ell k} = \max_{u \in \mathbf{U}} Y_{u\ell k} \in \{0, 1\}$. We implement two sampling strategies: a per-allele Gibbs sampler and a Metropolis-Hastings block move that updates all alleles of a single node jointly.

The Gibbs sampler draws a single allele $Y_{i\ell k}$ from its full conditional given the rest of $Y$, the observed data, and the model parameters. Because $Y_{i\ell k}$ is binary, the full conditional reduces to selecting the proposed state with probability

$$\frac{P(X \mid Y^{\text{prop}})P(Y^{\text{prop}})}{P(X \mid Y)P(Y) + P(X \mid Y^{\text{prop}})P(Y^{\text{prop}})}, \tag{101}$$

where $Y$ and $Y^{\text{prop}}$ differ only at $(i, \ell, k)$. Latent genetic state vectors are sparse, so a uniform random scan over alleles is inefficient. We instead use a non-uniform random scan that visits alleles discordant with the observed data more often, since the observed genetic state is highly informative of the latent state. Every allele retains positive selection probability, preserving irreducibility.

The Metropolis-Hastings sampler uses a proposal distribution that depends on the genetic states of parent infections, allowing sampling to be informed by both observed data and transmission network structure. We sample a parent set $\mathbf{U}$ uniformly from $\mathcal{U}_{i,\prec}$ and then draw $Y_i^{\text{prop}}$ from a kernel that conditions on $Y^{\mathbf{U}}$ and $X_i$. Marginalizing over $\mathbf{U}$ gives the proposal density:

$$g(Y_i^{\text{prop}} \mid X_i, Y, \mathcal{U}_{i,\prec}) = \frac{1}{|\mathcal{U}_{i,\prec}|} \sum_{\mathbf{U} \in \mathcal{U}_{i,\prec}} P(Y_i^{\text{prop}} \mid X_i, Y^{\mathbf{U}}), \tag{102}$$

where $P(Y_i^{\text{prop}} \mid X_i, Y^{\mathbf{U}})$ is the probability of drawing the allele collection $Y_i^{\text{prop}}$ given observed data $X_i$ and the parent-set indicator $Y^{\mathbf{U}}$.

We treat alleles as independent under the proposal:

$$P(Y_i^{\text{prop}} \mid X_i, Y^{\mathbf{U}}) = \prod_{\ell=1}^{L} \prod_{k=1}^{K_\ell} P(Y_{i\ell k}^{\text{prop}} \mid X_{i\ell k}, Y_{\ell k}^{\mathbf{U}}), \tag{103}$$

where $L$ is the number of loci and $K_\ell$ is the number of alleles at locus $\ell$. The independence assumption is a deliberate simplification of the proposal kernel; coupling across loci is handled by the joint conditioning on $\mathbf{U}$ and by the Metropolis-Hastings correction below.

The probability of proposing a single allele as present is

$$P(Y_{i\ell k}^{\text{prop}} = 1 \mid X_{i\ell k}, Y_{\ell k}^{\mathbf{U}}) = \begin{cases} 1 - \delta^+ & \text{if } Y_{\ell k}^{\mathbf{U}} = 1 \text{ and } X_{i\ell k} = 1, \\ \delta^- & \text{if } Y_{\ell k}^{\mathbf{U}} = 1 \text{ and } X_{i\ell k} = 0, \\ 0 & \text{if } Y_{\ell k}^{\mathbf{U}} = 0, \end{cases} \tag{104}$$

where $\delta^+, \delta^- \in (0,1)$ are user-defined tuning parameters of the proposal kernel. Their values play the role of false positive and false negative rates inside the proposal but are distinct from the observation-error parameters $\epsilon_i^+, \epsilon_i^-$ in Equation 92: $\delta^+, \delta^-$ control how aggressively the proposal flips alleles that are supported by the parent set, while $\epsilon_i^+, \epsilon_i^-$ define the likelihood of $X_i$ given $Y_i$. Any choice of $\delta^+, \delta^-$ yields a valid sampler because the Metropolis-Hastings ratio in Equation 106 corrects for the kernel. The third case forces $Y_{i\ell k}^{\text{prop}} = 0$ whenever no parent in $\mathbf{U}$ carries allele $k$, so the proposed latent state of node $i$ is contained in the parent-set indicator $Y^{\mathbf{U}}$.

We use this block move because the latent genetic state is sparse and strongly coupled across loci through the parent set, so jointly updating all alleles of node $i$ explores configurations that the per-allele Gibbs sampler reaches only one allele at a time. Writing $Y_i^{\text{prop}} \preceq Y^{\mathbf{U}}$ for the dominance relation $Y_{i\ell k}^{\text{prop}} \le Y_{\ell k}^{\mathbf{U}}$ at every $(\ell, k)$, the third case of Equation 104 sets $P(Y_i^{\text{prop}} \mid X_i, Y^{\mathbf{U}}) = 0$ whenever $Y_i^{\text{prop}} \not\preceq Y^{\mathbf{U}}$, so the marginal proposal in Equation 102 reduces to a sum over compatible parent sets:

$$g(Y_i^{\text{prop}} \mid X_i, Y, \mathcal{U}_{i,\prec}) = \frac{1}{|\mathcal{U}_{i,\prec}|} \sum_{\substack{\mathbf{U} \in \mathcal{U}_{i,\prec} \\ Y_i^{\text{prop}} \preceq Y^{\mathbf{U}}}} \prod_{\ell=1}^{L} \prod_{k=1}^{K_\ell} P(Y_{i\ell k}^{\text{prop}} \mid X_{i\ell k}, Y_{\ell k}^{\mathbf{U}}). \tag{105}$$

Adding or removing a single parent updates $Y^{\mathbf{U}}$ by a bitwise operation on the per-parent indicators. We cache each $Y_u$ with $u \in \mathbf{U}$, together with the per-allele factors of Equation 104 and refresh $Y^{\mathbf{U}}$ incrementally as $\mathbf{U}$ varies, which makes the sum tractable when $|\mathcal{U}_{i,\prec}|$ is large but candidate parent sets share many parents.

Each $Y^{\mathbf{U}}$ with $\mathbf{U} \in \mathcal{U}_{i,\prec}$ is a function of the latent states $\{Y_j : j \neq i\}$ alone, because the topological ordering excludes $i$ from $\mathbf{U}$. Replacing $Y_i$ with $Y_i^{\text{prop}}$ therefore leaves every $Y^{\mathbf{U}}$ fixed, so $g(Y_i \mid X_i, Y^{\text{prop}}, \mathcal{U}_{i,\prec}) = g(Y_i \mid X_i, Y, \mathcal{U}_{i,\prec})$, and the Metropolis-Hastings acceptance probability is

$$A(Y, Y^{\text{prop}}) = \min\left(1, \frac{P(X \mid Y^{\text{prop}}) P(Y^{\text{prop}})}{P(X \mid Y) P(Y)} \cdot \frac{g(Y_i \mid X_i, Y, \mathcal{U}_{i,\prec})}{g(Y_i^{\text{prop}} \mid X_i, Y, \mathcal{U}_{i,\prec})}\right), \tag{106}$$

where $Y^{\text{prop}}$ is the latent state obtained from $Y$ by replacing $Y_i$ with $Y_i^{\text{prop}}$ and both proposal densities are evaluated through Equation 105. The proposal is supported on $\{Y_i^{\text{prop}} : Y_i^{\text{prop}} \preceq Y^{\mathbf{U}}$ for some $\mathbf{U} \in \mathcal{U}_{i,\prec}\}$, so the reverse density vanishes and the move is rejected whenever the current $Y_i$ lies outside this support.

##### 1.11 Simulation Study Design and Procedures

###### 1.11.1 Simulation Procedure

We simulated data using a two-step process. First, we generated network topologies using a branching process combined with a merging step that adds superinfections to nodes, producing directed acyclic graphs (DAGs) with specific proportions of founders, average numbers of secondary cases ($R_0$), and proportions of superinfection as outlined in Table 2. Each network was simulated to have approximately 200 nodes. With the network topology fixed, we then simulated the transmission and observation processes.

| Setting | Values |
|---|---|
| **Founder Proportion** | 0.05, 0.2 |
| **Superinfected Proportion** | 0, 0.05, 0.1 |
| $\mathbf{R_0}$ | 0.5, 1, 1.5 |
| **Founder COI Rate** | 1, 2, 3 |
| **Total Network Topology Simulations:** | 54 |
| **Realizations per Topology:** | 10 |
| **Genotyping Panels per Realization:** | 3 |

**Supplementary Table 2:** Transmission network settings used in simulations. A total of 54 topologies were simulated, with each topology having 10 independent realizations of either the simplified model or the biologically based model. Each realization is characterized by 3 different genotyping panels, each with a different level of diversity.

For each simulated topology, we generated background allele frequencies from flat Dirichlet distributions of varying sizes, reflecting genetic loci with different diversity levels. We created three panels consisting of 50 allele frequency distributions of length 5, 10, or 20, representing genotyping panels of moderate, high, and very high diversity loci. Given these background population allele frequencies, we simulated the transmission process using two approaches: a simplified model matching our inferential framework and a biologically realistic model incorporating malaria parasite transmission biology. In both cases, we started with root nodes representing founder cases and simulated infection COI as draws from shifted Poisson distributions where the rate is the founder COI rate and the shift is 1 to ensure non-zero COI. We then sampled genetic states from background allele frequencies to produce infection genetic signatures. This process was repeated 10 times per topology to characterize variability in transmission network connectivity resolution due to varying information content of the genotyping panels.

###### 1.11.2 Simplified Model Transmission Simulation

In our simplified model, we simulated child infection genetics by sampling alleles from parent infection genetic states as described in subsection 1.6. This simplified approach may allow certain allele combinations that would be impossible in the biologically based model due to constraints requiring transmitted parasite genetic signatures to be consistent with single parent infections.

###### 1.11.3 Biologically Based Transmission Simulation

We simulated the transmission process sequentially from founder cases by modeling gametocyte transmission from each parent infection. Random gametocyte pairs form oocysts that undergo sexual recombination, producing sporozoites. The total number of oocysts was sampled from a zero-truncated negative binomial distribution with mean 2.7. We assumed genetic loci independence and that oocysts within a mosquito form from gametocytes from a single parent. We simulated sporozoite inoculation and establishment of child infections from each parent, with sporozoite numbers sampled from a lognormal distribution (mean 1.8, standard deviation 0.8). The genetic composition of sporozoite populations was uniform and evenly distributed from developed oocysts. For each individual, we simulated infection start times as uniform draws from the interval between parent infection and detection. Infection-to-detection periods were sampled from symptomatic and asymptomatic infection status distributions described in Equation 86, calculated with respect to final infections. Infection symptomatic status was randomly sampled with 0.5 probability of being asymptomatic.

###### 1.11.4 Observation Process Simulation

We simulated observed genetic states by masking underlying infection genetic signatures into binary vectors. Each allele at locus $\ell$ was randomly set or unset with false positive and false negative probabilities of $\frac{0.01}{K_\ell}$ and $\frac{0.1}{K_\ell}$ respectively, where $K_\ell$ is the total number of alleles at locus $\ell$.

###### 1.11.5 Model Fitting

As discussed in subsubsection 1.9.3, we applied prespecified constraints over possible parent sets by calculating pairwise genetic similarity metrics. For simulations, we used average identity by descent (IBD) calculated by Dcifer [7] as our genetic similarity metric. IBD provides a measure of genetic similarity explained by recent common ancestry rather than chance similarity relative to the parasite population. After calculating estimated IBD for each infection pair, we set a threshold of 0.1 on the confidence interval lower bound as our inclusion criterion for parent sets. This allows arbitrary numbers of infections in parent sets while removing highly unlikely parents without sufficient genetic similarity. We then fit the model to generate posterior distributions of parent set probabilities, the fundamental unit for calculating metrics of interest outlined in subsection 1.1.

###### 1.11.6 Case omission for robustness fits

Beyond the primary fit that conditions on every simulated infection in a replicate, we obtained a matched secondary fit by drawing a simple random 50% of infections to retain and removing the complementary infections from the analyzed DAG together with every transmission edge incident on a removed node. Amplicon observations and latent genetic updates were restricted to the retained infections. Dcifer pairwise IBD scores and the parent-set inclusion screen were recomputed on this censored graph using the same lower-bound threshold (0.1) as for the fully observed analysis. Edge-ranking diagnostics evaluate posterior and Dcifer scores on the censored graph (Figure 4, Figure 5). For recovery of average out-degree and of the fraction of infections whose true parent set includes an unsampled or external source, posterior expectations were computed on the censored graph but compared to targets taken from the corresponding full-network realization for the same replicate, so horizontal axes in the censored functional-recovery plots still reference the complete simulated cohort (Figure 6, Figure 7).

### 2 Supplementary Figures

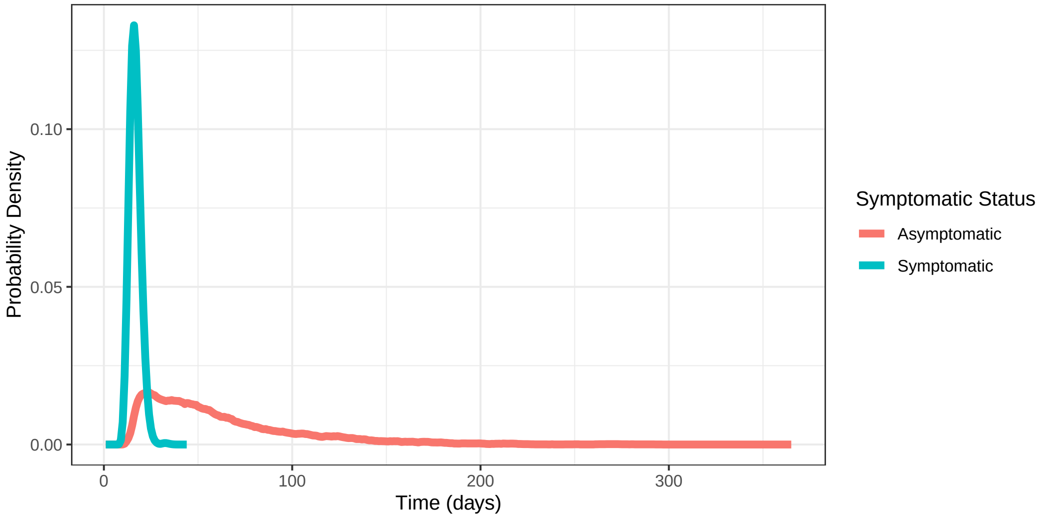

**Supplementary Figure 1: Probability density functions of the estimated IDP distribution for asymptomatic and symptomatic infections.** [4]

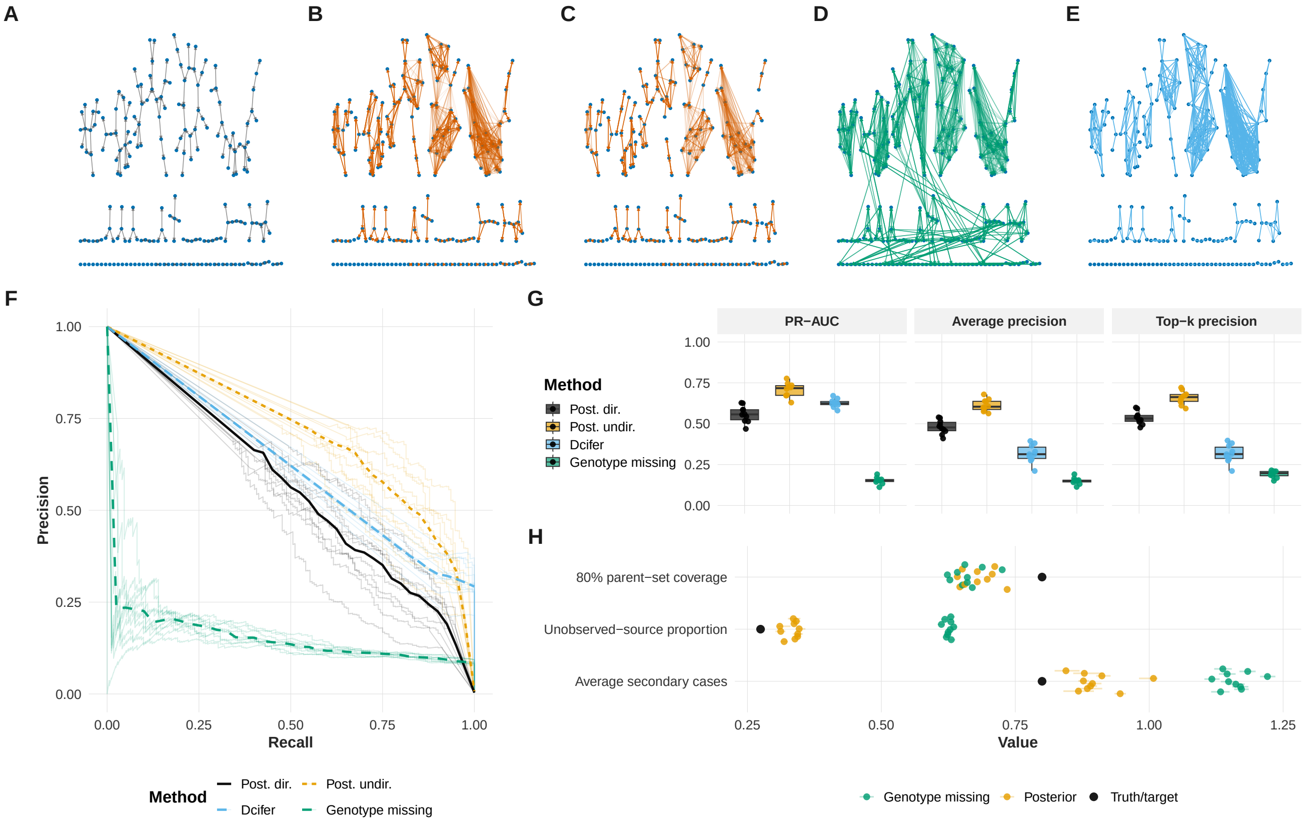

**Supplementary Figure 2: Scenario-group network diagnostics (biologically informed simulator).** One representative scenario group (matched simulation factors and genotyping panel tier, ten stochastic realizations; full observation). Top row, left to right: **A**, true directed transmission network among analyzed infections; **B**, posterior marginal directed edge probabilities (edge width and colour encode posterior support); **C**, directed edges in an 80% marginal credible set; **D**, marginal edge probabilities under genotype-missing fits (null genetic likelihood, with network priors and Dcifer-based parent-set restrictions retained); **E**, Dcifer undirected graph retaining pairs whose score exceeds the 0.9 quantile of strictly positive pairwise scores on the representative run. Bottom row: **F**, precision–recall style curves with scoring methods overlaid for the group; **G**, distributions of selected edge or ranking metrics across seeds; **H**, scalar functional summaries aggregated over the same realizations (e.g. average out-degree and unobserved-source summaries).

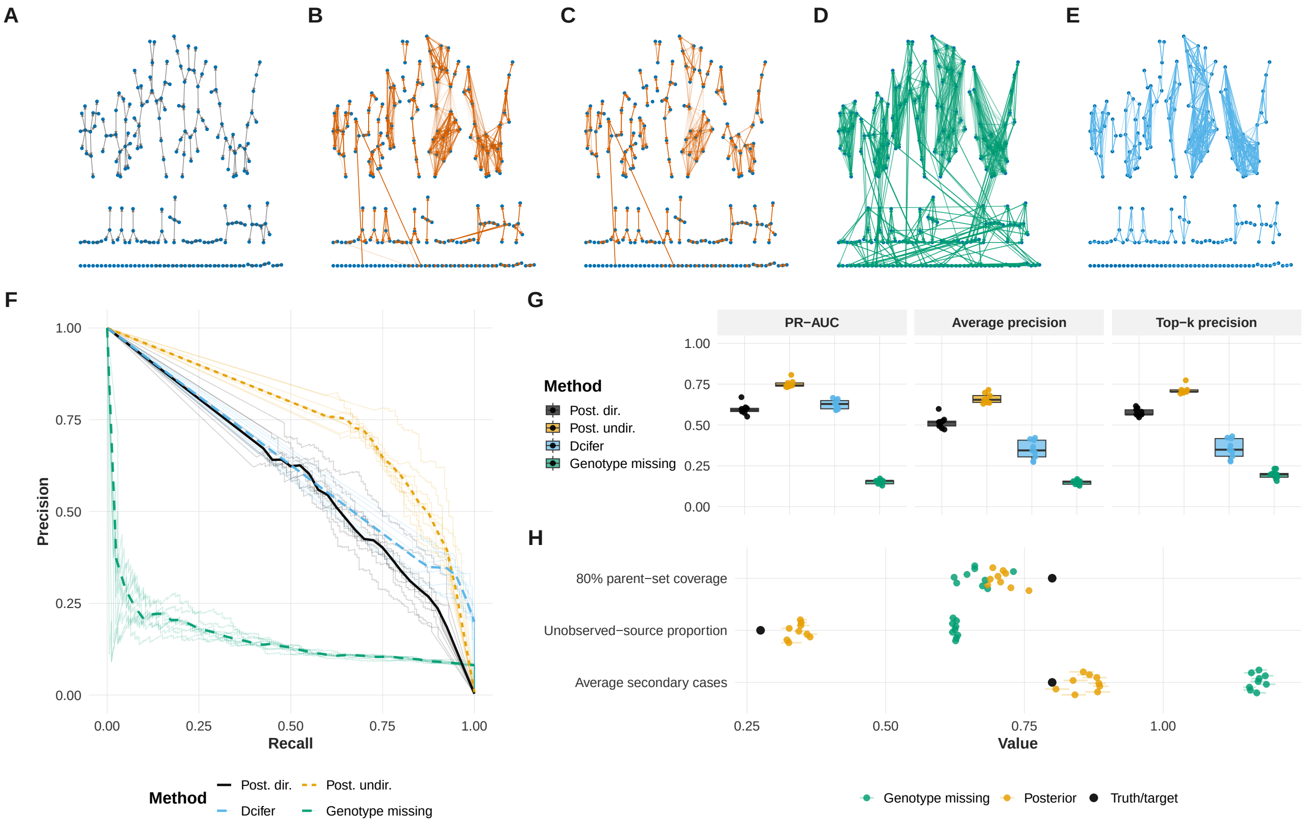

**Supplementary Figure 3: Scenario-group network diagnostics (simplified simulator matched to the inferential likelihood).** Same layout as Figure 2.

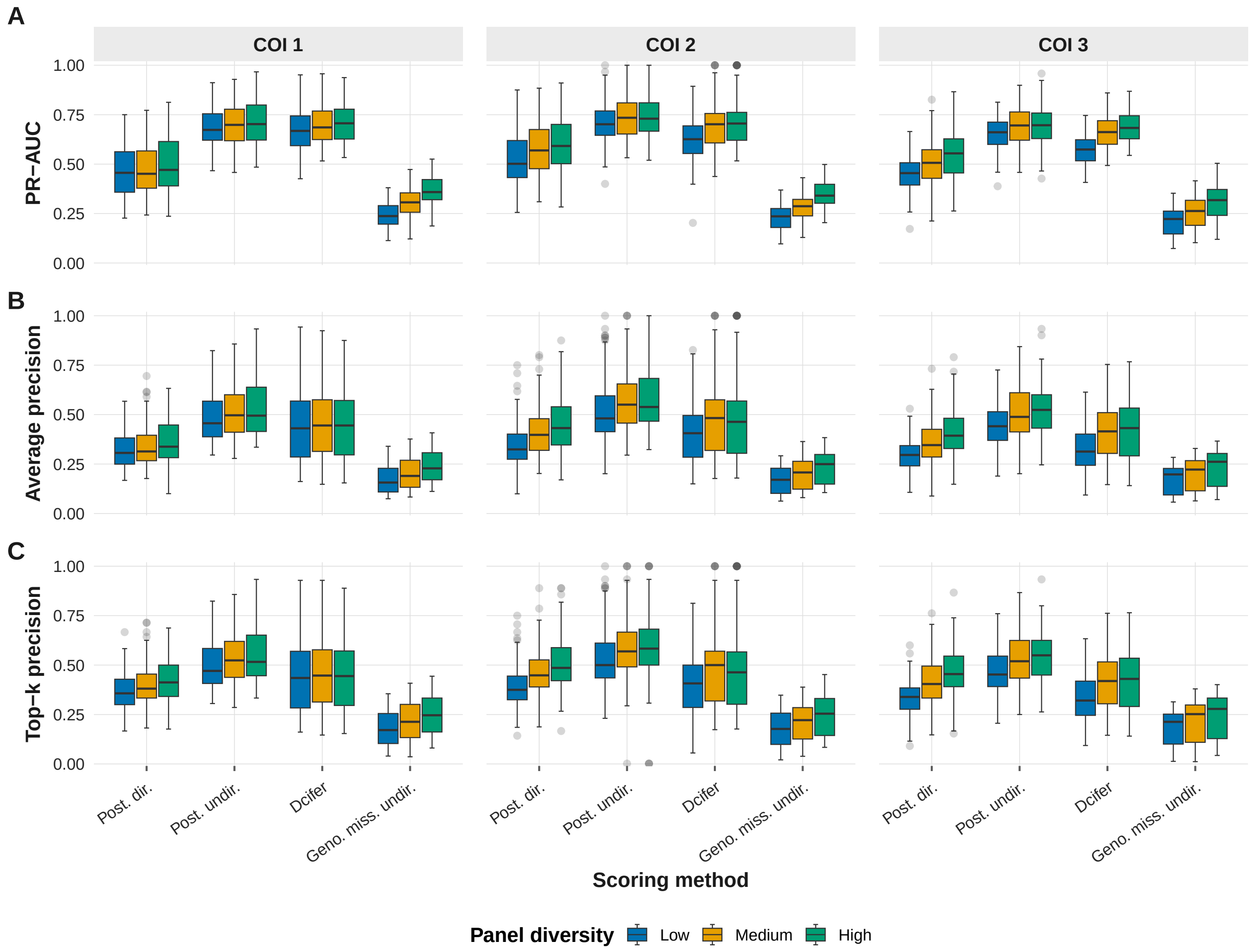

**Supplementary Figure 4: Edge classification metrics when half of infections are censored from the fitted graph (biologically informed simulator).** Each column is a founder COI stratum; colors indicate genotyping panel diversity. Rows show PR-AUC (panel A), average precision (B), and top-$k$ precision (C). Within each facet, boxplots contrast four replicate-matched scoring pipelines in fixed horizontal order: posterior directed rankings against directed truth; posterior undirected and Dcifer undirected rankings against symmetrized truth; and genotype-missing undirected rankings from fits that drop the genetic likelihood while retaining the same network priors and Dcifer-based parent-set restrictions. Posterior and Dcifer scores are computed on the 50% censored network for each run.

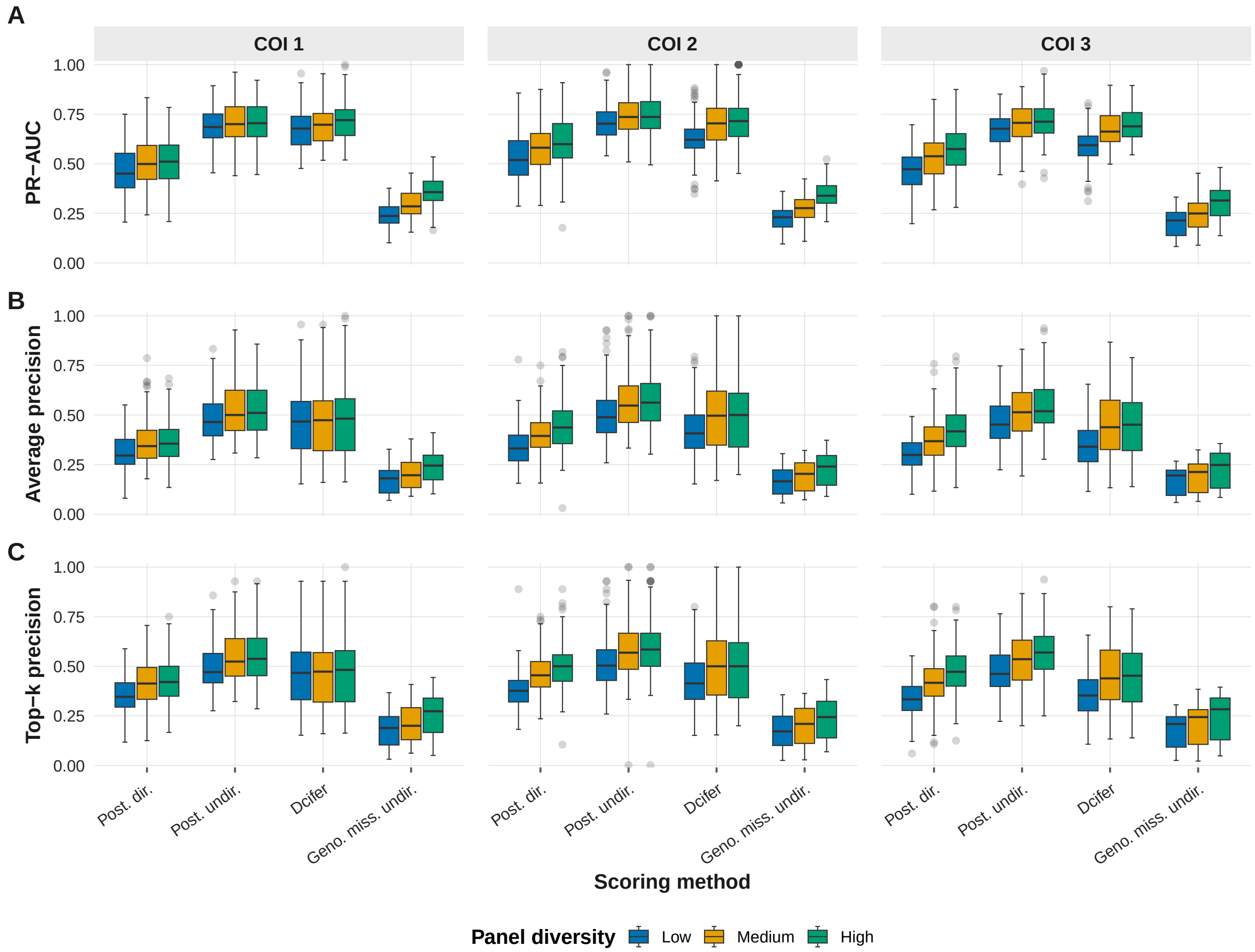

**Supplementary Figure 5: Edge classification metrics when half of infections are censored from the fitted graph (simplified simulator).** Same layout as Figure 4.

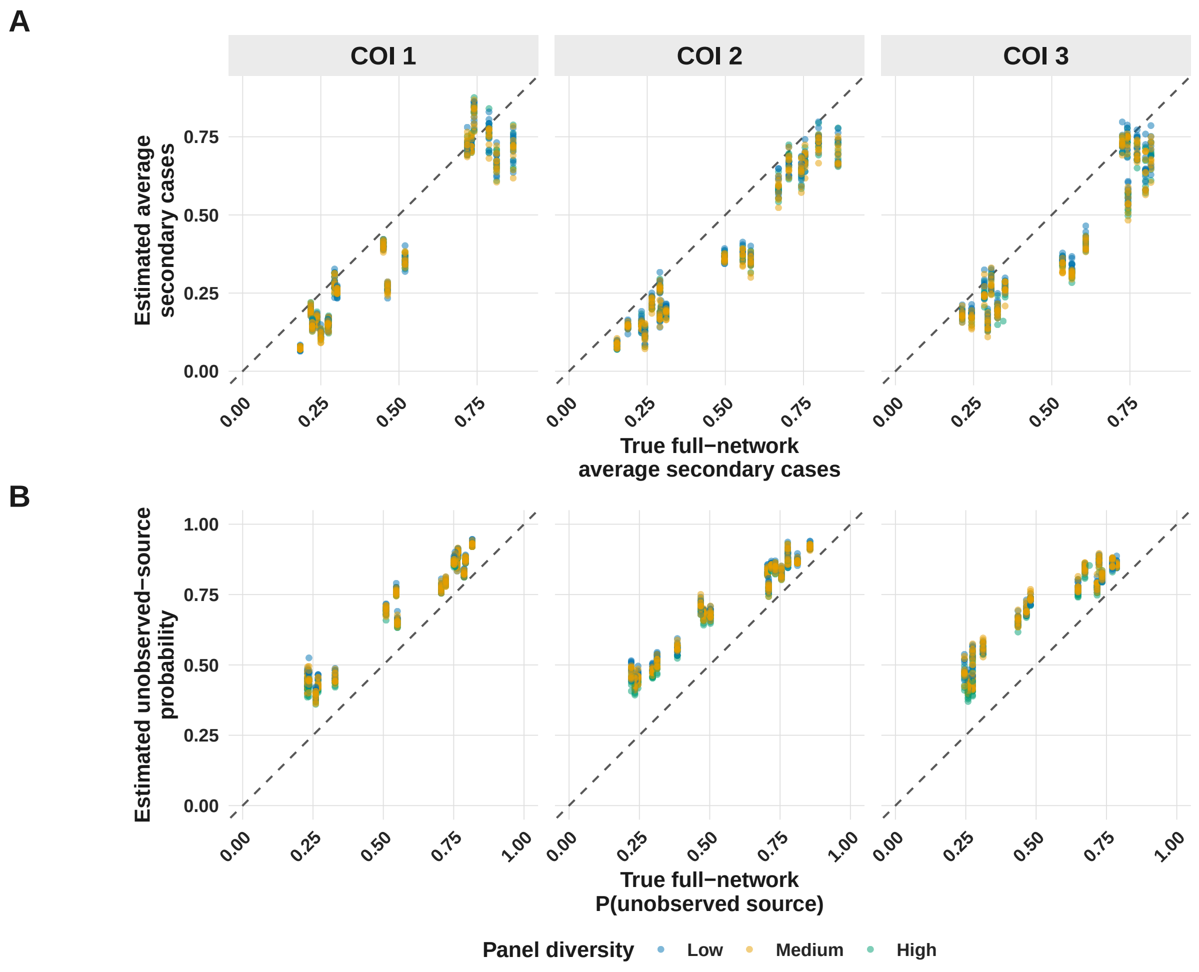

**Supplementary Figure 6: Recovery of average secondary cases and unobserved-source inclusion under 50% case censoring (biologically informed simulator).** Posterior summaries are computed on the 50% censored observed subset for each replicate. Panel A compares true full-network average out-degree to its posterior expectation under censoring. Panel B compares the simulated full-network fraction of infections whose parent set includes an unsampled or external source to the mean posterior unobserved-source inclusion probability under censoring. Columns stratify by founder COI; colors indicate genotyping panel diversity.

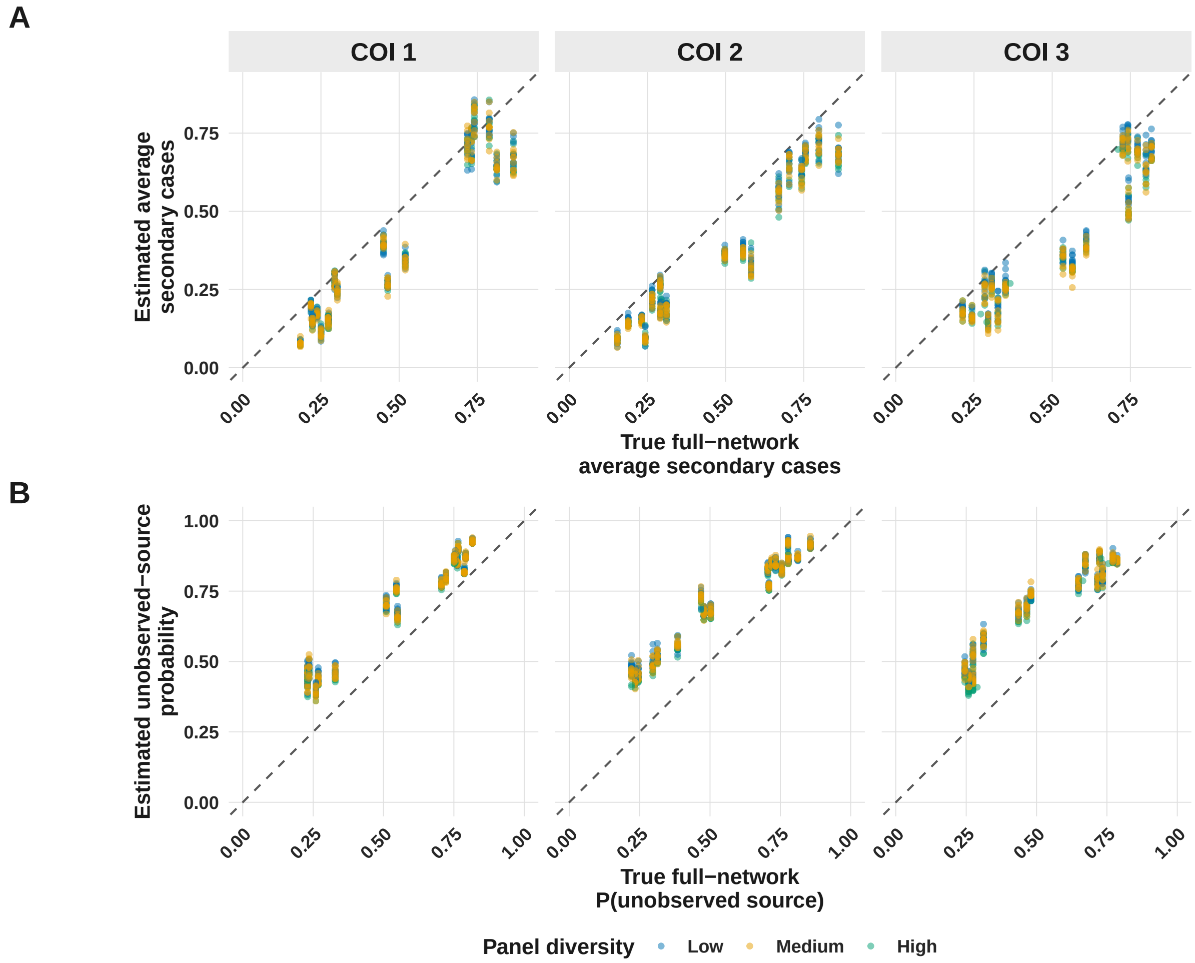

**Supplementary Figure 7: Recovery of average secondary cases and unobserved-source inclusion under 50% case censoring (simplified simulator).** Same layout as Figure 6.

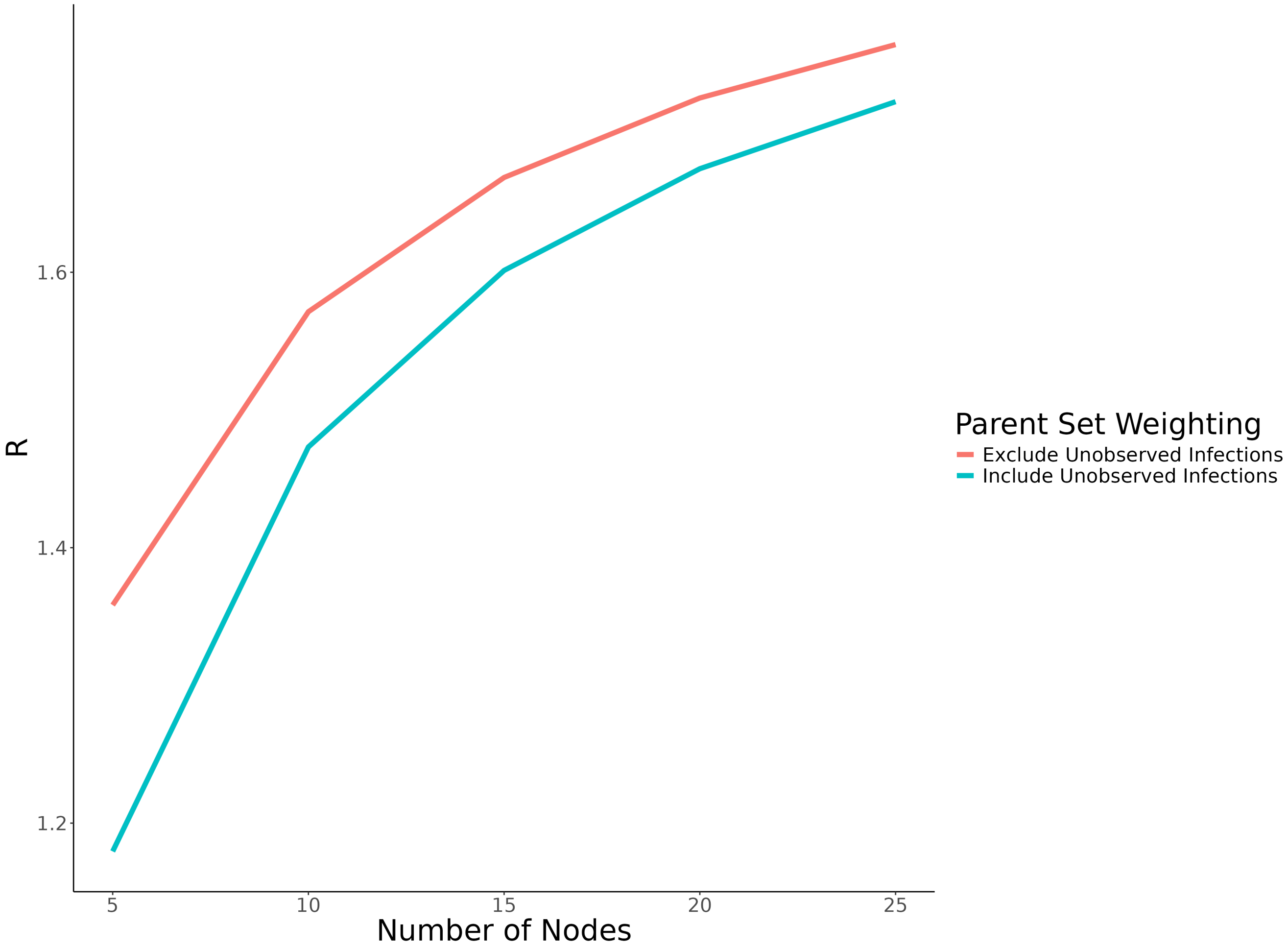

**Supplementary Figure 8: Exact R values for different parent set weighting schemes across a range of network sizes under a fixed constraint on parent sets to include up to 2 observed parents and 1 unobserved parent.** Excluding unobserved infections from possible parent sets results in a higher R value under a fixed topological ordering when weighting all parent sets equally. This situation may arise to some degree when genetic data supports connectivity between infections, but does not differentiate between infections, such as a cluster of genetically identical infections. In contrast, including unobserved infections in possible parent sets results in a lower R value under a fixed topological ordering. This situation would arise when there is insufficient genetic data to differentiate between local transmission and introduction or importation, such as when genetic marker diversity is very low and infections are genetically identical.
